# Mitigation of Antimicrobial Resistance Genes in Greywater Treated at Household Level

**DOI:** 10.1101/2023.02.15.528737

**Authors:** Weitao Shuai, Daniella Itzhari, Zeev Ronen, Erica M. Hartmann

## Abstract

Greywater often contains microorganisms carrying antimicrobial resistance genes (ARGs). Reuse of greywater thus potentially facilitates the enrichment and spread of multidrug resistance, posing a possible hazard for communities that use it. As water reuse becomes increasingly necessary, it is imperative to determine how greywater treatment impacts ARGs. In this study, we characterize ARG patterns in greywater microbial communities before and after treatment by a recirculating vertical flow constructed wetland (RVFCW). This greywater recycling method has been adopted by some small communities and households for greywater treatment; however, its ability to remove ARGs is unknown. We examined the taxonomic and ARG compositions of microbial communities in raw and treated greywater from five households using shotgun metagenomic sequencing. Total ARGs decreased in abundance and diversity in greywater treated by the RVFCW. In parallel, the microbial communities decreased in similarity in treated greywater. Potentially pathogenic bacteria associated with antimicrobial resistance and mobile genetic elements were detected in both raw and treated water, with a decreasing trend after treatment. This study indicates that RVFCW systems have the potential to mitigate antimicrobial resistance-related hazards when reusing treated greywater, but further measures need to be taken regarding persistent mobile ARGs and potential pathogens.

## 1. Introduction

As freshwater is increasingly limited and prone to anthropogenic pollution, wastewater treatment facilities play an important role in reuse of valuable water. With proper treatment, recycling greywater (all non-toilet wastewater resulting from households) can increase the efficiency of water usage and benefit arid regions.^1^ Israel is a semi-arid country and the supply of potable water relies mainly on groundwater, surface water and desalination of seawater.^2^ Because of this limitation, some small communities and households in Israel have adopted recirculating vertical flow constructed wetlands (RVFCW) for on-site greywater treatment for local landscaping irrigation.^3, 4^ While greywater reuse is important for addressing water scarcity, greywater contains both bacteria and micropollutants, making it a meeting point of human-related microorganisms and disinfectants, biocides, personal care products, and pharmaceuticals. Exposure of microorganisms to antimicrobials or biocides at environmental concentrations can enhance the spread of antimicrobial resistance in microbial communities.^5, 6^ The co-occurrence of bacteria and micropollutants in greywater potentially promotes the discharge of antimicrobial-resistant bacteria (ARB) and dissemination of antimicrobial resistance genes (ARG). Exposure to ARB and ARGs therefore poses a potential health hazard to household and community members. The discharge of ARB and ARGs from “hot spots” like water treatment facilities is an essential part of the antimicrobial resistance spread in the environment. Therefore, characterization of the antimicrobial resistance in these systems is prioritized in order to better assess the health risks and develop policies.^7^

Previous studies have reported occurrence of antibiotics, herbicides^8^, ARB and ARGs^9-11^ in off-grid greywater reuse systems. However, these studies focused on a limited number of ARB and ARGs using culture-based and qPCR-based analyses. The current study aims to give a more holistic overview of the taxonomic profile and ARG composition in both raw and treated greywater by examining the metagenome. In so doing, we reveal potential selective pressures at play in these engineered systems and substantiate the needs of monitoring the performance of decentralized water reclaim systems like RVFCWs to better mitigate risk of ARB and ARG spread.

## 2. Methods and materials

### 2.1 Description of the RVFCW

The greywater treatment system studied in this research is the RVFCW described by Gross *et al*^4^. The RVFCW consists of two 500 L containers stacked vertically. Influent from individual household, raw greywater is introduced into the surface of the upper container, and then trickles through the passively aerated wetland bed to enter the lower container reservoir. The greywater is recirculated from the reservoir back to the upper container wetland bed and then pumped out with a hydraulic retention time of 6-8 hours. All the sampled households use the same water source consisting of a mixture of desalinated seawater and groundwater. During the sampling period, the source water chlorine concentration was between 0.3-0.5 mg/L, turbidity was between 0.3 and 0.7 NTU, and *E. coli* was always less than 1 CFU per 100 ml (https://mywater.health.gov.il/WaterNetwork). The recycled greywater is used for landscaping irrigation on the premises, and it can pose health hazards to humans via direct contact and indirectly via contact with irrigated soil.^12^ The five RVFCW facilities in this study are highly similar except for variation in filter bedding and the presence of vegetation (Table 1).

**Table 1.**
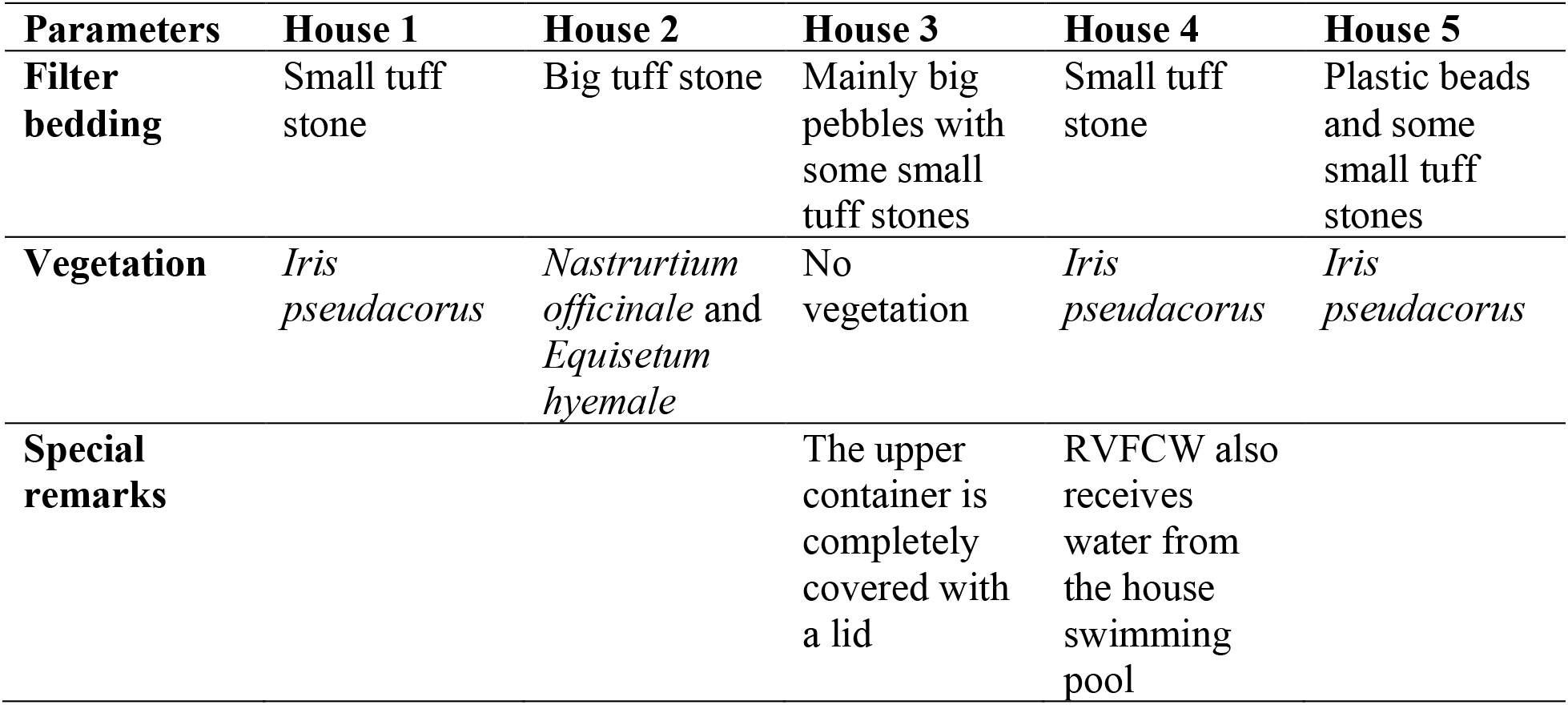
Configuration and operating conditions of the RVFCW facilities.

| Parameters | House 1 | House 2 | House 3 | House 4 | House 5 |
| --- | --- | --- | --- | --- | --- |
| **Filter bedding** | Small tuff stone | Big tuff stone | Mainly big pebbles with some small tuff stones | Small tuff stone | Plastic beads and some small tuff stones |
| **Vegetation** | *Iris pseudacorus* | *Nastrurtium officinale* and *Equisetum hyemale* | No vegetation | *Iris pseudacorus* | *Iris pseudacorus* |
| **Special remarks** |  |  | The upper container is completely covered with a lid | RVFCW also receives water from the house swimming pool |  |

### 2.2 Sample description and quantitative PCR assay

Raw and treated greywater samples were taken in October 2020 from each household and transported to the laboratory within 4 hours of sampling. Aliquots of 150 mL (300 mL for house 3) raw greywater and 1 L treated greywater were filtered through a glass fiber filter (type A/E 46mm PALL) and 0.2 µm filter (PALL). DNA was extracted using a DNeasy PowerWater Kit (Qiagen, product code 14900-100-NF) for raw greywater samples and MasterPure™ Complete DNA and RNA Purification Kit (Lucigen, product code MC85200) for treated greywater samples. Quantification of the abundance of bacterial genes was performed using quantitative PCR (qPCR). qPCR was performed with 20 µl reaction volumes composed of qPCRBIO SyGreen Blue Mix Lo-ROX (PCR Biosystems). Primer set 341F/518R (5’-CCTACGGGAGGCAGCAG-3’/5’-ATTACCGCGGCTGCTGG-3’, annealing temperature 55 ºC) targeting the 16S rRNA gene was used for total bacteria quantification.^13^ Calculation of the complete gene copies of target genes was based on known copies of a standard reference plasmid. Copies of the 16S rRNA gene were normalized by volumes of raw or treated greywater samples filtered for DNA extraction. Standard curve, calculated efficiency, and reaction protocol of qPCR experiments are available in Supporting Information (Figure S9 and Figure S10).

### 2.3 Metagenomic sequencing analysis

Sequencing libraries were constructed using an Illumina DNA Prep (previously known as Nextera DNA Flex Library Prep) Kit and paired-end 150-bp shotgun metagenomic DNA sequencing was performed using an Illumina HiSeq 4000 at the NUSeq Core Facility at Northwestern University. Two kit controls (Qiagen glass filter and PALL; Lucigen glass filter and PALL) were also sequenced to reveal contaminants. The Qiagen glass filter and PALL kit control generated 4704 forward and reverse reads, while Lucigen glass filter and PALL kit control generated 14890 forward and reverse reads. Greywater sequencing samples were decontaminated using KneadData v0.7.4 (https://github.com/biobakery/kneaddata) to eliminate human genome contamination and sequences from kit control samples using default settings. MultiQC v1.10.1^14^ was used to check sequence quality of all greywater samples before and after decontamination. Metagenomic coverage and sequence diversity were estimated using Nonpareil 3 (v3.304)^15^ on the decontaminated, paired sequence output from KneadData. Short read-based taxonomic classification was performed using Metaxa2 (v2.2)^16^ and MetaPhlAn 2.0.^17^ Since Metaxa2 uses the small subunit (SSU; 16S/18S) rRNA gene and MetaPhlAn 2.0 uses clade-specific markers, Metaxa2 tends to recognize more taxa from poorly characterized environmental communities, whereas MetaPhlAn 2.0 tends to be more accurate at lower taxonomic levels. Therefore, the Metaxa2 output was used for illustrating the overall taxonomic diversity, and MetaPhlAn 2.0 was used for identifying the presence of potential pathogens.

Short reads were assembled using metaSPAdes from SPAdes v3.15.3 release.^18^ QUAST 5.0.2^19^ was used for assembly quality assessment. Assembly-based taxonomic classification was performed on the resulting scaffolds using Kraken 2 with the default database.^20^ Antibiotic resistance genes were detected using both short read sequences and assembled scaffolds using Resistance Gene Identifier (RGI version 5.2.1; https://github.com/arpcard/rgi) from the Comprehensive Antibiotic Resistance Database (CARD) using database version 3.1.2.^21^ RGI bwt mode (short read-based analysis) gives mapped read counts, which were converted to the relative abundance of each ARG allele in unit of reads per kilobase per million mapped reads (RPKM) for comparison between samples. For the RGI bwt output, mapped reads with MAPQ ≥ 50, coverage ≥ 90% of the reference length or read length (150 bp) were attained for downstream analysis. The RGI main program was applied on assembled scaffolds (contigs) to explore the co-occurrence of specific genes and taxa. For RGI main output, identity ≥ 95%, best hit bit score ≥ 50 and percentage length of reference sequence between 90% and 110% were applied to obtain high-quality hits. Mobile genetic elements (MGE) were identified from the assemblies using MobileElementFinder^22^ and outputs were filtered by identity ≥ 95% and coverage ≥ 90%. UpSet plots of ARG and MEG were generated using ComplexHeatmap R pakage version 2.10.0^23^.

### 2.4 Statistical analysis

Principle component analysis (PCA) was conducted to examine the taxonomic and ARG composition. Permutational multivariate analysis of variance (PERMANOVA; number of permutations = 9999) was conducted to identify important factors explaining the variance. Python package scikit-learn (0.22.1)^24^ and R package VEGAN (2.4-2)^25^ were used for statistical analyses.

## 3. Results and discussion

### 3.1 Overview of metagenomic sequencing data

The absolute 16S rRNA gene copy number did not change significantly after the greywater treatment (Figure 1; two-tailed, paired *t*-test, p > 0.05). After decontaminating the raw metagenomic sequences, the 10 greywater samples yielded 29.5 million paired-end reads on average with over 20 million reads for all samples (Sequencing and decontamination results presented in Table S1). The metagenomic coverage estimated by Nonpareil 3 had a relatively large range and a negative relationship with the sequence diversity (Figure S2). Nonpareil applies a redundancy-based approach to provide an abundance-weighted coverage of the metagenome, which represents the fraction (range 0-1) of the microbial community sampled by DNA sequencing. The difference in metagenomic coverage between raw and treated greywater samples was more pronounced than the difference between households. Treated greywater sequences have distinctly higher diversity (two-tailed, paired t-test, p < 0.05) and lower coverage (two-tailed, paired t-test, p < 0.05) compared to raw greywater sequences. Metagenomic assemblies from treated greywater samples on average have more contigs, but smaller N50 value compared to the assemblies from raw greywater samples (two-tailed, paired *t*-test, p < 0.05; Table S2). The higher sequence diversity and lower metagenomic coverage values of treated greywater sequences may have resulted in more fragmented contigs and lower genome fractions (two-tailed, paired t-test, p < 0.05; Table S2) in the metagenome assemblies. Metagenomics has enormous potential to facilitate monitoring of diverse pathogens and ARGs, but it still has conceptual and technical limits of detection and quantification in the practice of environmental surveillance^26^, including insufficient coverage exacerbated by community complexity. Therefore, comparisons between raw and treated greywater are mostly based on short read-based results in this study, whereas assembly-based results are viewed as qualitative supporting information.

**Figure 1.**
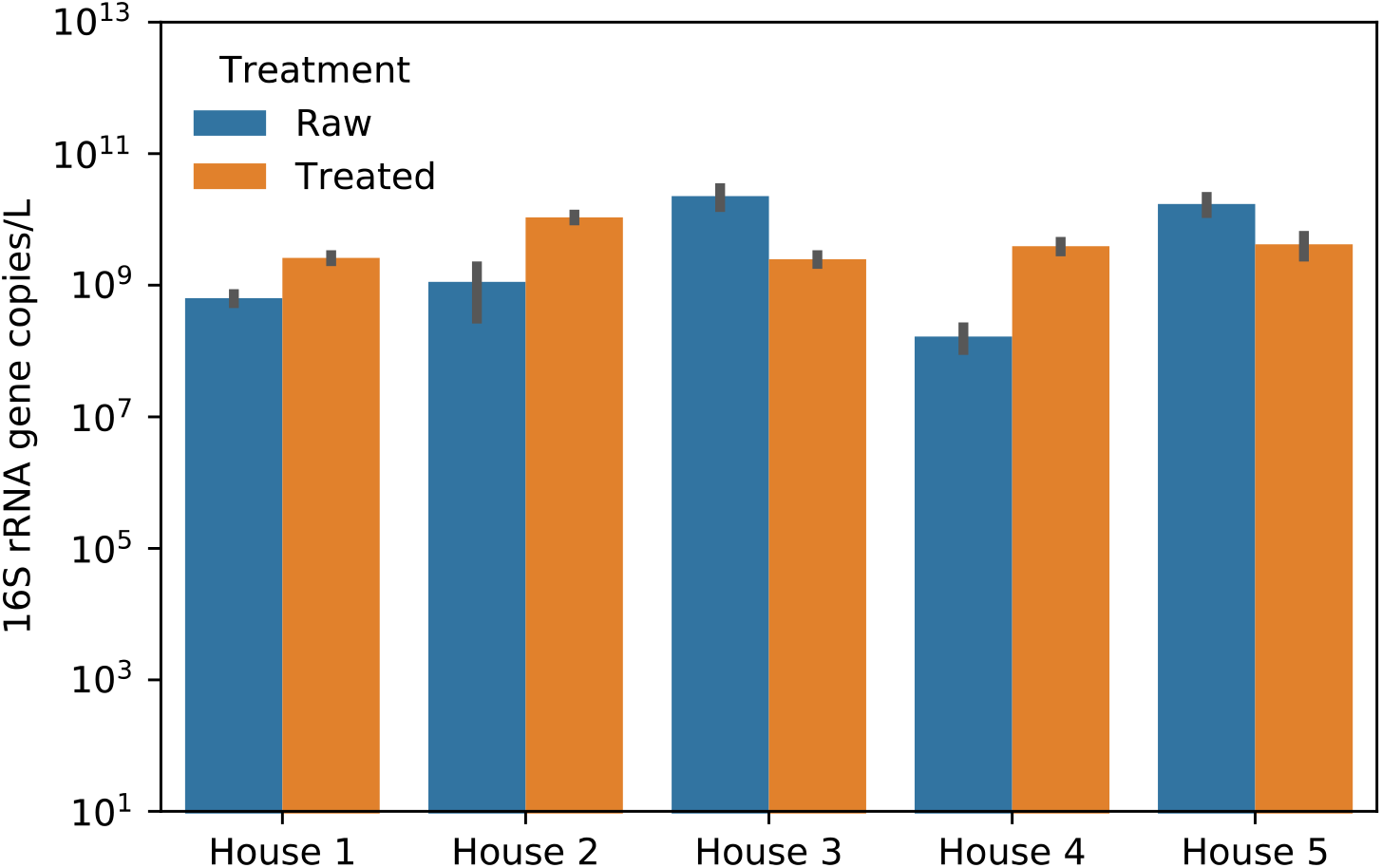
16S rRNA gene quantification in five houses before and after treatment by RVFCW (Error bars are standard deviations of qPCR technical replicates, n=3).

### 3.2 Overall microbial communities in different greywater treatment facilities diverge after treatment

Proteobateria was the predominant classified phylum among the greywater samples, followed by Bateroidetes and Firmicutes (Figure S2). These three phyla comprised more than 50% of the microbial community in all samples. The high relative abundance of Chromalveolata, Metazoa, Alveolata and Rhizaria in treated greywater samples could be due to the presence of vegetation and various filter bed materials in the RVFCW, which provided habitat for these eukaryotes.

The greywater samples showed diverse taxonomic profiles at the genus level, with no universally dominant genera ranked by their relative abundance among all 10 samples, within either raw or treated groups (Figure 2A). The microbial community structures of greywater samples from different houses were more similar to each other before treatment than after treatment, as evidenced by PCA (Figure 2B). PERMANOVA results show that different houses had larger effect size (21.7% of the total sum-of-square, p < 0.05) compared to treatment (14.4% of the total sum-of-square with borderline statistical significance p = 0.08). The fact that the five studied houses are receiving the same water source could contribute to the similarities of microbial communities in the raw greywater samples, although differences in characteristics like occupancy, personal care product usage, water usage habits would shape the microbial community of greywater at each household. The difference in the microbial and chemical species input as well as the slight difference in RVFCW parameters can further influence the microbial community structure as the greywater gets recirculated in RVFCW for hours. The distinct environmental factors at each household can contribute to the divergence of greywater microbial communities since contact time is extended during the treatment process compared to merely flowing through the household water pipes.

**Figure 2.**
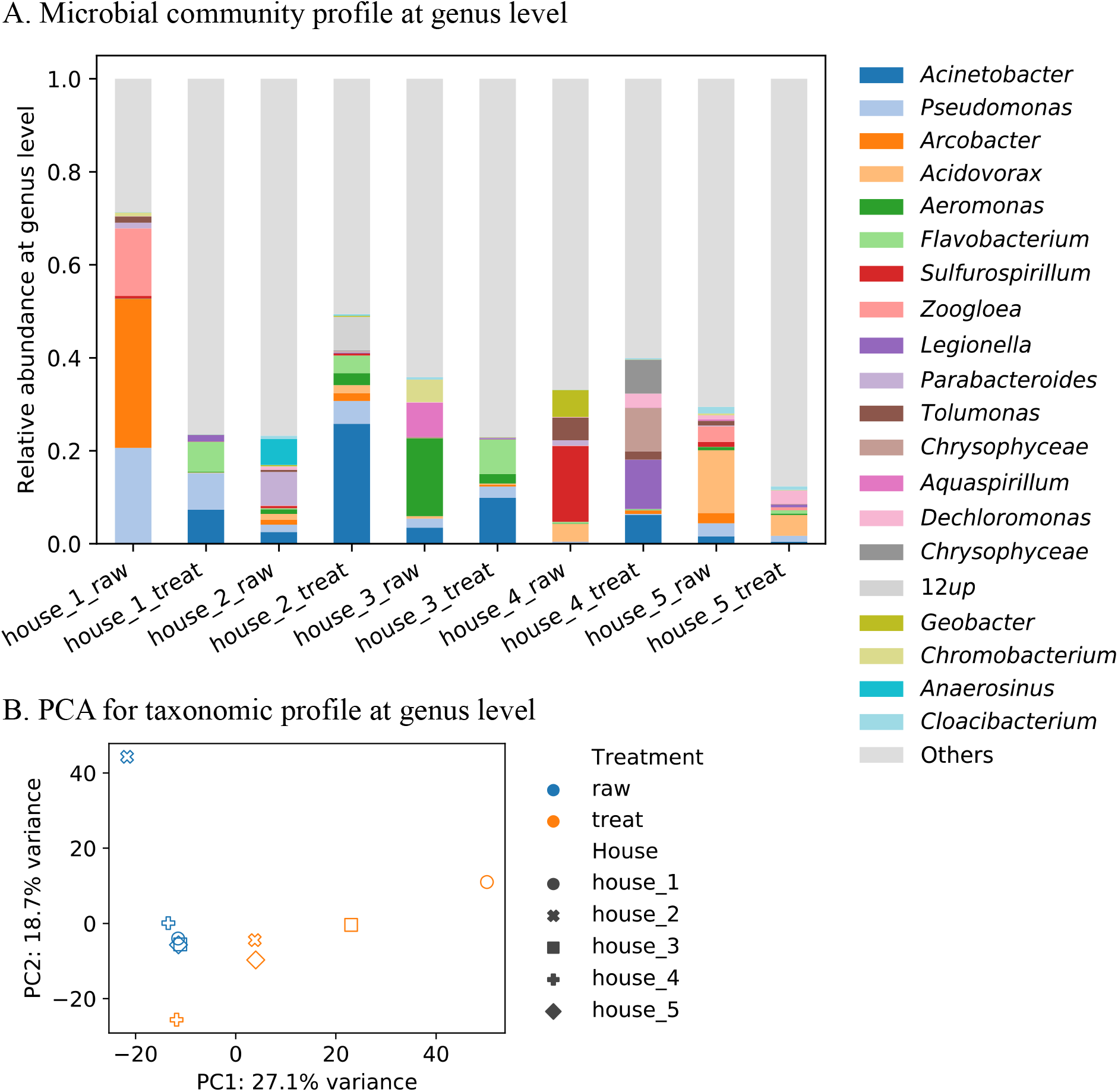
Taxonomy profiles of all greywater samples based on Metaxa2 output. (A) Top 20 genera ranked by average relative abundance across all greywater samples. Unidentified taxa at different classification levels other than genus are incorporated in “Others”. (B) Principal component analysis (PCA) for taxonomic profile at genus level.

While no single genus dominated multiple samples, a considerable number of genera that dominated individual samples contain common pathogenic species, namely *Acinetobacter, Pseudomonas, Arcobacter, Aeromonas, Legionella* and *Parabacteroides*. Similar ranking of potential human pathogens (*Acinetobacter, Pseudomonas, Aeromonas*, and *Legionella)* was also observed in a study on a wastewater treatment plant (WWTP) and its receiving waters.^27^ At finer taxonomic resolution, common human pathogens *Citrobacter freundii, Klesiella oxytoca*, and *Desulfovibrio desulfuricans* were found in the top 50 most abundant species identified by signature sequences using MetaPhlAn 2.0, along with some less common human pathogens which are possibly responsible for opportunistic infections in immunocompromised hosts (Table S3). The appearance of these human pathogens at such high relative abundance among the microbial communities in the greywater samples, especially in treated greywater samples, emphasizes the need for monitoring the potential of health hazards posed by on-site greywater reuse. We define a hazard potential which considers both the relative abundance of clinically important taxa and total biomass for each greywater sample. The hazard potential log_2_ fold change 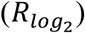 is calculated as:

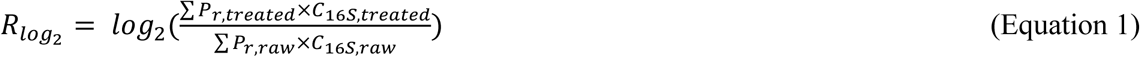

Where ∑*P*_*r*_ is sum of relative abundance of clinically important (“pathogen”) taxa identified in top 50 species detected by MetaPhlAn 2.0 (Table S3), *C*_16*S*_ is the concentration of 16S rRNA gene copies quantified by qPCR in greywater samples. Results showed that hazard from potential pathogens could even increase after treatment by RVFCW in some households due to an increase of clinically important taxa or an increase of total biomass (Figure 3, house 2 and 4). Therefore, it is important to understand the fate of potential human pathogens and ARG carriers in these decentralized water treatment systems, along with different environmental and operational factors, in order to provide recommendations for more efficient and effective water treatment to reduce the overall hazards. Busgang *et al*. assessed health risks associated with greywater reuse using a quantitative microbial risk-assessment (QMRA) model for some individual pathogens under different exposure scenarios and provided maximum concentrations of the pathogens required to keep the risk below the maximum acceptable level. ^12^ Accompanied with qPCR, metagenomic sequencing can provide useful information to feed the exposure assessment for multiple pathogens, determining the overall risk for all water-related illnesses in a holistic manner.

**Figure 3.**
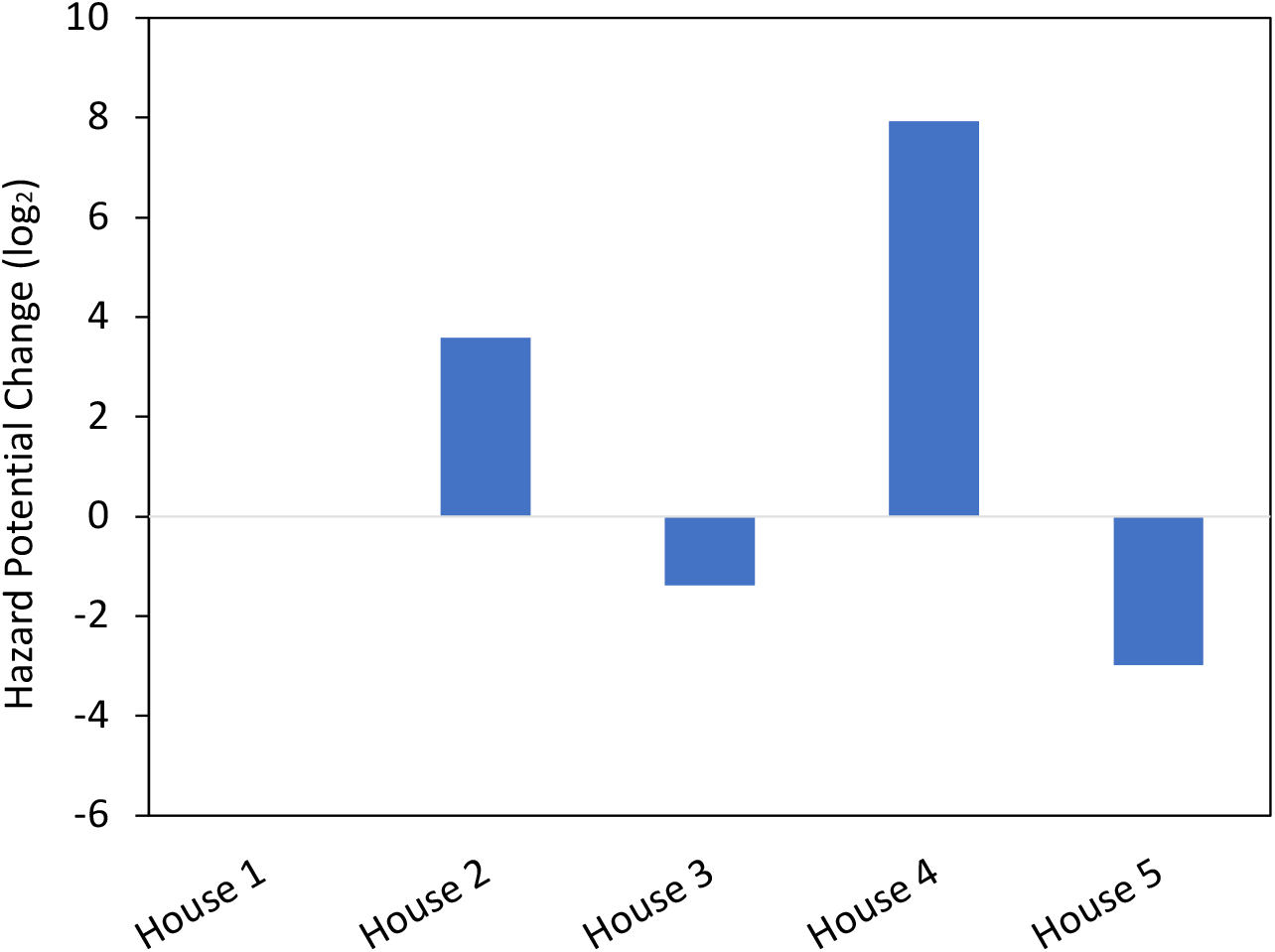
Hazard potential from clinically important taxa log_2_ fold change after treatment by RVFCW. Positive numbers indicate increased hazard potential after treatment.

### 3.3 ARGs are less abundant in treated greywater than in raw greywater

The number of distinct ARG alleles found in greywater samples decreased in post-treatment samples for all five houses (Figure 4A). Grouping the five households together, 71 and 37 different ARG alleles were identified in raw and treated greywater samples, respectively. The results are comparable to the numbers of ARGs found in municipal WWTP, with 6 or 19 different ARGs in two WWTP metagenomic projects^28^ and a total of 53 ARGs found among six water samples from farm and WWTP effluent.^29^ Cumulative relative abundances (RPKM) of ARG from the short read-based RGI output are higher in raw greywater samples than treated greywater samples (Figure 4B) except for house 2. The hazard potential of ARGs in greywater was calculated using Equation 1 where cumulative relative abundance of clinically important taxa was replaced by cumulative relative abundance of ARG (Figure 4C). Results showed that the hazard potential from ARGs could increase even when ARG relative abundance and diversity both decrease after treatment by RVFCW (house 1 and 4).

**Figure 4.**
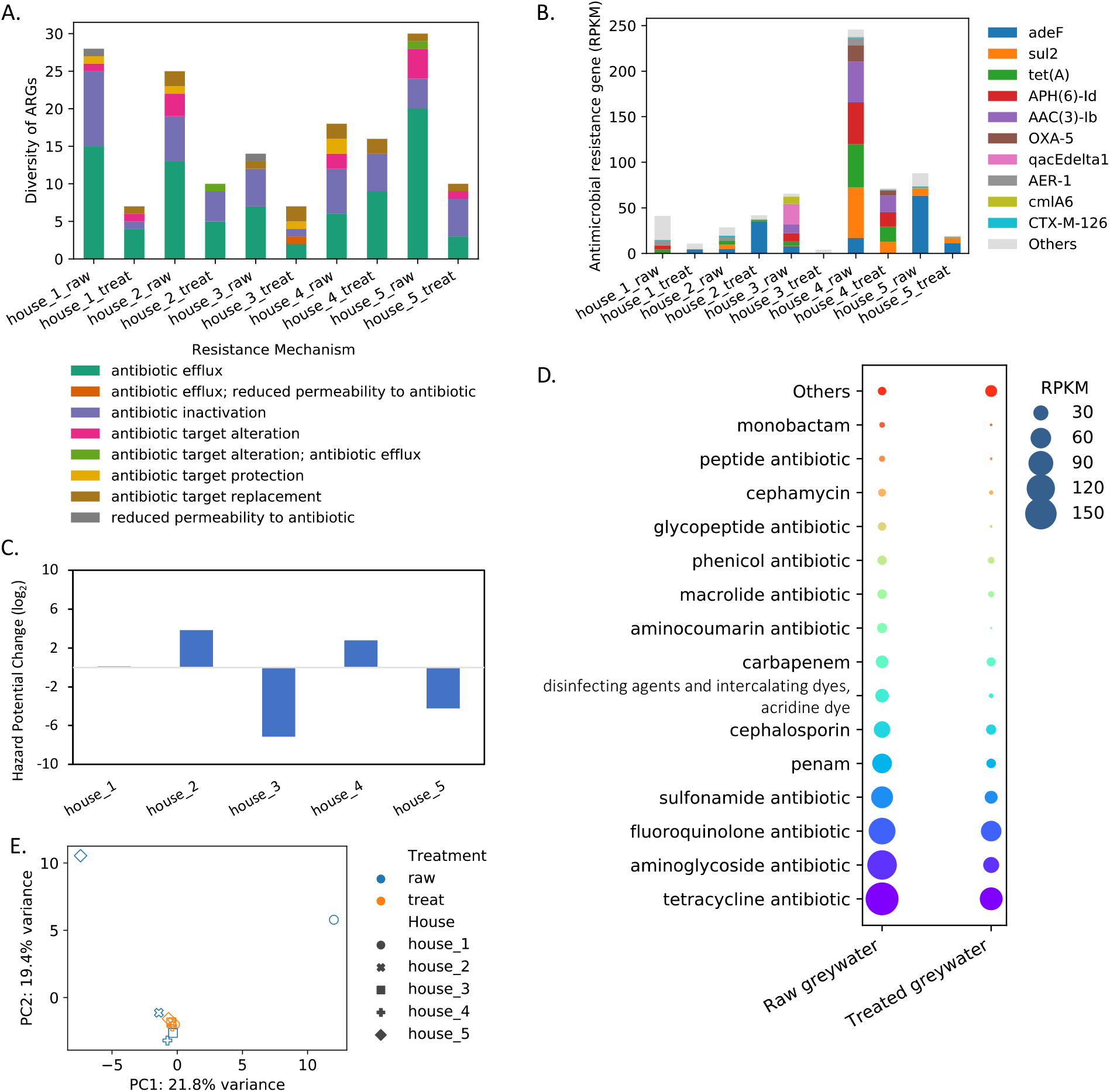
Antimicrobial resistance gene (ARG) profiles in greywater samples from short read-based analyses. (A) Richness grouped by resistance mechanism. (B) Relative abundance in RPKM. (C) Hazard potential log_2_ fold change after treatment. (D) Top 15 ARG drug classes ranked by their cumulative relative abundance. Note that many ARGs confer resistance to more than one drug class, thus their relative abundance was included in multiple drug classes. Different colors indicate different drug classes. (E) Principal component analysis for ARG profile.

Resistance genes involved in efflux comprised 12-96% of the ARG abundance for individual greywater samples, which summed up to 48% of the total ARG abundance for all (raw and treated) samples. The cumulative abundance of efflux pump genes decreased after treatment, with the exception of house 2 (10.7 and 40.5 RPKM in raw and treated samples, respectively). As stated by Martinez *et al*.,^30^ the upregulation, not the mere presence, of the efflux pump genes in a metagenome should be considered as real threat, as the wild-type version of these genes should not confer resistance after transfer to susceptible bacteria. Since DNA sequencing data from this current study can only illustrate gene presence, high efflux gene proportions among ARG are not necessarily a concern. The cumulative abundance of non-efflux-mediated resistance genes decreased by 36-98% after treatment in all five households (Figure S3), indicating reduced potential of ARG transfer to pathogenic or other bacteria. ARGs were grouped by the targeting drug class to infer overall resistance potential of raw *versus* treated greywater samples (Figure 4D). Relative abundance of all 15 top ARG groups decreased after treatment, which presented a more consistent trend than comparing the abundance of any individual ARG. The decrease of relative abundance of ARG-carrying taxa possibly contributed to the decrease of relative abundance of ARGs through treatment. Positive Spearman’s rank correlations (r ≥ 0.6, p < 0.05) can be observed for some taxa (genus level) and ARG alleles that appeared to have high abundance in greywater (Table S4). With correction of total biomass ratios, increased abundance after treatment by RVFCW can be observed mainly at house 2 and house 4 (Figure S4), which is consistent with the trend of hazard potential change for possible pathogens and ARGs.

### 3.4 Groups of ARGs presented in greywater samples at relatively high abundance

Out of the identified ARG alleles, only *adeF* (encoding the membrane fusion protein of the multidrug efflux complex AdeFGH) is shared among all five raw greywater samples. At the gene family level, only genes belonging to resistance-nodulation-cell division (RND) antibiotic efflux pump and major facilitator superfamily (MFS) antibiotic efflux pump are shared by all 10 greywater samples. This is not surprising as RND family transporters are the most commonly found bacterial efflux systems in Gram-negative organisms,^31^ and MFS is the second largest group of membrane proteins.^32^ ARGs conferring resistance to fluoroquinolone antibiotic drugs found in this study, which was the third most abundant group (Figure 4D), all belong to RND, MFS or ATP-binding cassette (ABC) antibiotic efflux pump gene families.

Sulfonamide resistance *sul* genes and beta-lactamase genes appear to be important in the greywater ARG composition, making up 14.6% and 9.6% of the total abundance for the 10 greywater samples respectively. Resistance genes grouped by drug class sulfonamide and beta-lactams (penam, cephalosporin, carbapenem, cephamycin, and monobactam), which were also identified in urban WWTPs,^33^ ranked high in the greywater samples (Figure 4D).

In addition, ARGs (*qacEΔ1, mdtM, mdtN, mdtO*, and *mexW*) with resistance to disinfecting agents and intercalating dyes (acridine dye) drug class were notable in raw greywater (Figure 4D). These genes all belong to RND or MFS antibiotic efflux pump gene family. Specifically, *qacEΔ1* (ranked the seventh most abundant ARG across the 10 greywater samples; Figure 4B) is recognized as an antiseptic-resistance rather than an antibiotic resistance gene that can be found in both Gram-negative and Gram-positive bacteria^34, 35^. As biocides (including disinfectants, sanitizing agents and chemical cleaning agents) are commonly present among other micropollutants in greywater^36^, their role in shaping the antimicrobial resistance profile should be considered. Unlike antibiotics, biocides usually interact nonspecifically on microbial targets. Intrinsic antimicrobial resistance mechanisms like active efflux and reduced uptake are usually not specific to substrates either,^37^ meaning they could be part of the natural bacterial response to stress or injuries imposed by biocides.^38^ The causative role of biocides exposure in antimicrobial resistance promotion is yet to be confirmed,^39^ although some studies have shown positive relationships^40, 41^ even in drinking water systems.^42^

### 3.5 Environmental factors at the households shaped the distinct ARG compositions

Unlike the taxonomic profiles, the ARG profiles of the treated greywater samples are more similar to each other compared to those of the raw greywater samples (Figure 4E). This divergence of taxonomic profiles and convergence of ARG profiles could be related to the increase of taxonomic diversity (Figure S5) and decrease of ARG diversity (Figure 4A) of treated greywater compared to raw greywater. PERMANOVA results showed that neither different houses nor treatment had a statistically significant (p > 0.1) effect size on the variance of ARG abundance, which can be attributed to the small sample size and scarcity of shared ARG alleles among samples (Figure S6). The majority of ARG alleles are unique to each sample, with 74% of the ARGs only detected in one individual sample. Thus, the ARG composition is possibly highly dependent on distinctive environmental factors at each household rather than the water supply. For comparison, a study on promotion of antimicrobial resistance by disinfectants in drinking water systems suggested that the precise antimicrobial resistance traits detected in each system varied significantly, and bacterial community structure was the strongest determinant of antibiotic resistance ontology composition.^42^ This could be true in our system as well, since taxonomic profiles were more divergent (Figure 2B) and diverse (Figure S5) after treatment where shared ARG alleles were fewer (Figure S6). When total biomass (as 16S rRNA gene copies) was taken into account, the increased biomass after RVFCW treatment at house 2 and house 4 acted as an important driver for a positive change of hazard potential for both possible pathogens and ARG alleles (Figure 3 and Figure 4).

High levels of surface-active material, micropollutants, and cleaning products which are ubiquitous in raw greywater can co-select for ARGs.^43-46^ Greywater treatment operational conditions can also be contributors. For example, a main operation parameter of the system in removing ARG is the recirculation flow rate (RFR), which influences the efficiency of both chemical and biological contaminant removal. It was shown that increasing RFR from 2.5 m^3^ h^-1^ to 4.5 m^3^ h^1^ substantially improved the quality of the treated effluent.^47^ It was also shown that low oxygen in oxygen-based membrane biofilm reactor led to the accumulation of biofilm that increased the level of ARGs in greywater.^48^ The filter bed of RVFCW can accumulate biofilm; however, it is not stressed by oxygen in our system because the recirculation constantly aerates the filter. A study compared three different constructed wetland configurations for greywater pathogen removal and found that the aerobic unsaturated (vertical flow) wetland was the most suitable technology among the three configurations based on pathogen enumeration.^49^

In our study, interesting relationships were observed between distinct conditions at individual household and its ARG composition. House 4 had a considerably higher cumulative ARG abundance comparing to other houses, as the untreated greywater contained much higher abundances of *sul2, tetA, aph(6)-Id*, and *aac(3)-Ib* genes (Figure 4B). These genes contributed largely to the top ranked resistance drug classes in greywater samples, namely tetracycline antibiotic, aminoglycoside antibiotic, and sulfonamide antibiotic (Figure 4D). The major difference between house 4 and other houses was that the RVFCW also received swimming pool effluent. The concentration of chlorine inside the pool ranged between 1-3 ppm, and 500 liters of this water was emptied every month to the greywater. Chlorine disinfection was reported to enrich ARGs *ermB, tetA, tetB, tetC, sul1, sul*2, *sul3, ampC, aph(2’)-Id, katG*, and *vanA* in a full-scale wastewater treatment plant^50^, *ampC, aphA2, bla*_*TEM-1*_, *tetA, tetG, ermA*, and *ermB* in a drinking water plant^51^ where the chlorination time was 30 min to 4 hours in these systems. Karumathil *et al*.^52^ reported that antibiotic resistance genes in *Acinetobacter baumannii* were up-regulated upon exposure to 2 ppm chlorine. Based on evidence shown in this study and available literature, it is possible that untreated greywater microbial community from house 4 was affected by chlorine exposure, thus the functional potential of antibiotic resistance was enriched. The ARG composition of the untreated house 4 greywater agrees with enrichment of ARG groups induced by chlorination. Nevertheless, richness and cumulative ARG abundance decreased after treatment (Figure 4AB), indicating that RVFCW could partially counteract the potential adverse impacts posed by application of chlorine disinfectants although potential hazard was driven up by increased total biomass after treatment.

In this current study, greywater from five houses had unique ARG compositions even though they shared same water supply and had similar microbial community structure before treatment. Treatment at RVFCWs could have exacerbated the slight differences in operational and environmental factors at each house, resulting in more divergent taxonomic profiles (Figure 2B) and unique ARG compositions (Figure S6). Although it is desirable to estimate an ARG or ARB removal efficiency for decentralized water reclamation systems to facilitate microbial risk assessment in regard to antimicrobial resistance spread, varied operational and environmental factors at individual facilities complicate the outcome and make it hard to generalize design and regulatory recommendations. This is a major challenge for accurate quantitative microbial risk assessment (QMRA) on single organisms, as model inputs like log reduction values and horizontal gene transfer rate are dependent on treatment conditions.^53, 54^ Nevertheless, metagenomic sequencing can be a valuable monitoring tool for non-target, broad screening of ARGs, ARB, and MGEs in the QMRA hazard identification framework^55^ due to its ability to identify ARB that are not detected by culture-based methods and higher efficiency of identifying ARGs compared to PCR-based methods.

### 3.6 Associations between ARGs and microbial taxa in raw and treated greywater

Among the top 10 most abundant ARGs detected in the greywater samples (Figure 4A), *tetA, aph(6)-Id*, and *qacEΔ1* genes and their variants are reported to be observed in *Escherichia coli* according to the curated resistance gene database CARD.^21^ However, *E. coli* was not identified in read-based taxonomic classification. Instead, unclassified *Escherichia* (or unclassified *Escherichia-Shigella*) was found but not at high relative abundance. The short read-based taxonomic classification may have limitations in identifying some species, leading to under-estimation of species that are associated with antibiotic resistance. Therefore, assembly-based analysis was conducted to better characterize the relationship between taxa and ARGs.

To identify the association of ARGs and taxa in the greywater microbial community, the assemblies (contigs) containing ARGs were classified using Kraken 2 (Figure S7). There were 120 counts of ARGs identified on contigs in raw greywater samples and 66 counts in treated greywater samples. Among these ARG-containing contigs, 87 and 44 counts had taxonomic classification down to species level. This reduced frequency of ARG-species association after treatment by RVFCW is consistent with the short read-based results for ARG abundance and diversity discussed above.

The most frequent taxon-ARG pair is *Pseudomonas alcaligenes* – *rsmA* in raw greywater and *Acinetobacter gyllenbergii* – *adeF* in treated greywater. *P. alcaligenes* is a rare opportunistic human pathogen^56^ and *A. gyllenbergii* has been isolated from human clinical specimens^57^; both are reported to exhibit resistance to multiple antibiotics.^56, 58^ *Pseudomonas aeruginosa, E. coli, Citrobacter freundii, P. alcaligenes*, and *Klebsiella pneumoniae* are the top five ARG-associated species in raw greywater samples (Figure S7A), while *Klebsiella grimontii, Acinetobacter baumannii, A. gyllenbergii, E. coli*, and *Acinetobacter disperses* are the top five ARG-associated species in treated greywater samples (Figure S7B). This result aligns with the taxonomic profiles in which *Acinetobacter* and *Pseudomonas* are the two most abundant genera identified by Metaxa2 (Figure 2), and *C. freundii* and *P. alcaligenes* are among the top 50 species identified by MetaPhlAn 2.0 (Table S3). Although both pathogenic and nonpathogenic bacteria can display antimicrobial resistance as their natural response to antibiotics^59^, the top ranked species associated with ARGs are common or opportunistic human pathogens in our system. *P. aeruginosa, E. coli*, and *C. freundii* showed association with more than five ARGs in raw greywater samples. After treatment, *P. aeruginosa* and *C. freundii* were not in the top five ARG-associated species (Table S5).

Brown *et al*. found that patterns of ARG contextualization differed between short-read/hybrid assemblies and long-read assembly and suggested that adequate short-read sequencing depth was essential to harness the full potential of the proposed hybrid assembly approach.^60^ In our study, the higher taxonomic diversity and lower metagenome coverage of treated greywater samples (Figure S1) may affect the overall quality of assemblies, resulting in an observed decrease in occurrence of taxon-ARG associations detected in treated greywater samples compared to raw greywater samples (Figure S7, Table S5). Nevertheless, as the trend of ARG occurrence agrees with the short-read based analysis, we believe that the assembly-based ARG results reflect a change of ARG composition and provide information on the relationships between specific ARGs and species. This association between pathogenic microorganisms and ARGs again urges us to continue monitoring and assessment of how decentralized water treatment facilities perform from the perspective of mitigating antimicrobial resistance dissemination.

### 3.7 Potential mobility of ARGs in greywater samples decreased after treatment

Gene mobility is an import evaluative factor when it comes to assessing the health risk of ARGs^61^, since mobile ARGs promote dissemination of resistance among previously susceptible microorganisms. We further characterized the ARG composition in greywater based on their potential mobility using CARD prevalence database (version 3.0.9). ARGs that have non-zero prevalence in NCBI plasmids comprised of 29 – 77% of the total ARG diversities in the greywater samples, and their diversities before and after treatment also followed the overall trend of total ARGs diversity change (Figure 4A; Figure 5A). Although the diversity of ARGs observed in the plasmid database decreased after greywater treatment, it is worth noting that these ARGs are abundant in the total ARG composition, and the proportions remained high in four out of five houses even after treatment by the RVFCW facilities (Figure 5B). Interestingly, house 3 has the only facility without vegetation and the upper container is completely covered with a lid (lower dissolved oxygen level is expected), which is also the only facility that had substantial decrease in the proportion of plasmid-associated ARGs. This contradicted with other studies where low oxygen in membrane biofilm reactor increased the level of ARGs in greywater^48^ and vegetated constructed wetlands removed significant number of antibiotic-resistant bacteria than non-vegetated constructed wetlands from hospital wastewater.^62^ It is hard to conclude from this single example from our study the role of RVFCW operating conditions (vegetation, lid application, limitation of atmospheric deposition, flow configurations) in the treatment of particular groups of ARGs. However, with the RVFCW’s potential of removing pathogens and reducing ARG spread, it is worth investigating the relationships between RVFCW operating parameters and efficiencies of ARG and pathogen removal with detailed survey and monitoring in the future.

**Figure 5.**
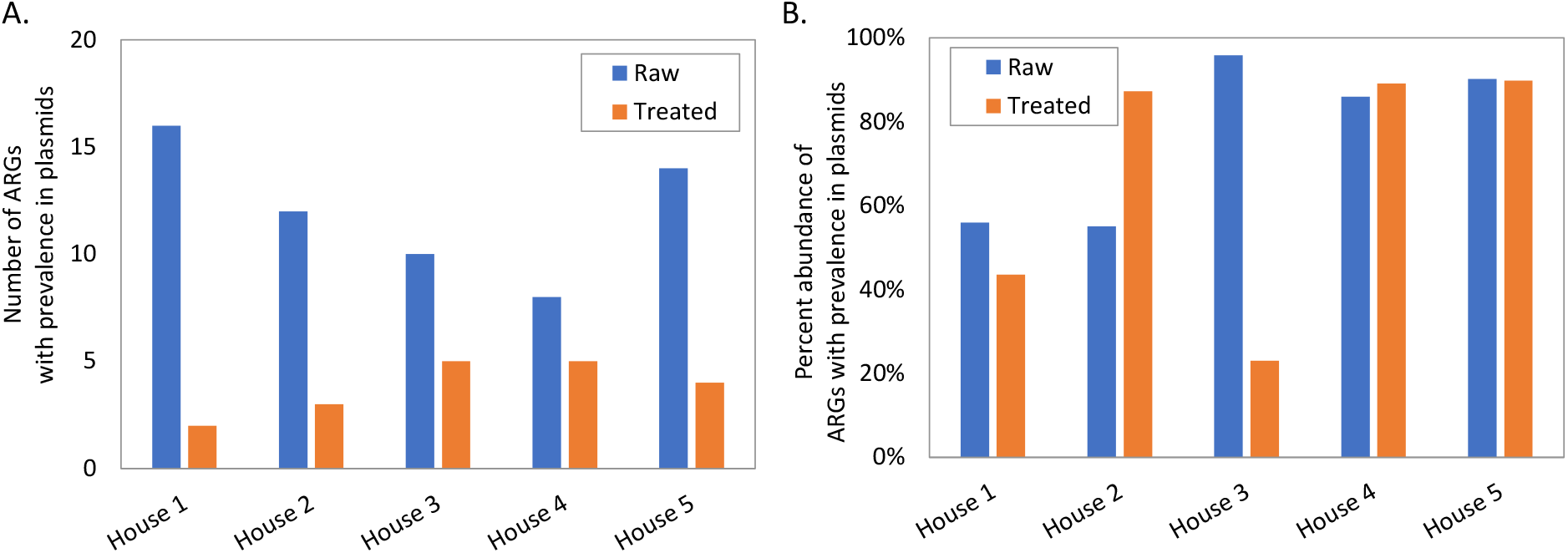
Number of ARGs with non-zero prevalence in NCBI plasmids (A) and their percent abundance of the total ARG abundance (B). Results are from short read-based ARG abundance analysis.

Mobile genetic elements (MGEs) detected by MobileElementFinder using the assemblies presented similar trend: fewer MGEs were found in treated than raw greywater samples (Figure S8A) with the exception of house 4, which also had the smallest decrease in richness of ARGs (Figure 4A). The majority (90 - 100%) of detected MGEs were insertion sequences. Similar to ARGs found in the greywater samples, MGEs appeared to be unique to individual samples instead of shared between treatment condition or household (Figure S8BC). The top five MGE-associated species in our greywater samples were *Aeromonas caviae, Acinetobacter junii, P. aeruginosa, Acinetobacter johnsonii*, and *Aeromonas hydrophila*, which are common, opportunistic or rare human pathogens.^63-66^ The co-existence of MGE and ARG on the same contig was very rare in our greywater samples, potentially due to insufficient sequencing depth. Therefore, we do not attempt to link the presence of MGEs to specific ARG alleles. Although results from assemblies are more qualitative than quantitative in this study, we observed a decrease of occurrence and diversity for both ARG-pathogen pairs and MGEs after greywater treatment in the studied household-level RVFCW facilities.

## Conclusion

The greywater metagenomes from five households in Israel allow us to investigate the change in microbial community and ARG composition before and after treatment by RVFCW facilities. Human pathogens and other microorganisms associated with ARGs, which are not are not detected by culture-based methods, were detected in both raw and treated greywater samples. While taxonomic diversity increased after treatment, the diversity and abundance of ARGs, MGEs, and ARG-associated species decreased after treatment. Results from this study indicate that treatment of raw greywater by RVFCW can reduce the potential of ARG and antibiotic resistance pathogen dissemination. Nevertheless, we are fully aware of the limitations of this study such as insufficient sequencing depth, lack of greywater micropollutant monitoring, and small sample numbers. However, interesting observations from this study provide information that can direct future investigations on RVFCWs in Israel as well as other similar decentralized greywater treatment systems. Our study demonstrated that there are needs for more in depth monitoring of decentralized water treatment facilities like the greywater recycling systems on the antimicrobial resistance front. Further efforts are needed to more clearly discern factors promoting the persistence of ARGs and MGEs specifically to mitigate the potential health risk.

## Acknowledgement

Authors would like to acknowledge The Zuckerberg Institute for Water Research-Northwestern University Program for financial support in this project. This research was supported in part through an award from the 2020 NUSeq Pilot Project Program sponsored by Illumina, as well as through the computational resources and staff contributions provided for the Quest high performance computing facility at Northwestern University which is jointly supported by the Office of the Provost, the Office for Research, and Northwestern University Information Technology. We thank Alexander G. McFarland for help in troubleshooting metagenomic analysis tools, En-ling Wu for help in clinically important species identification, and Stefanie Huttelmaier and Jiaxian Shen for help in internal review of the manuscript.

## For Table of Contents Only

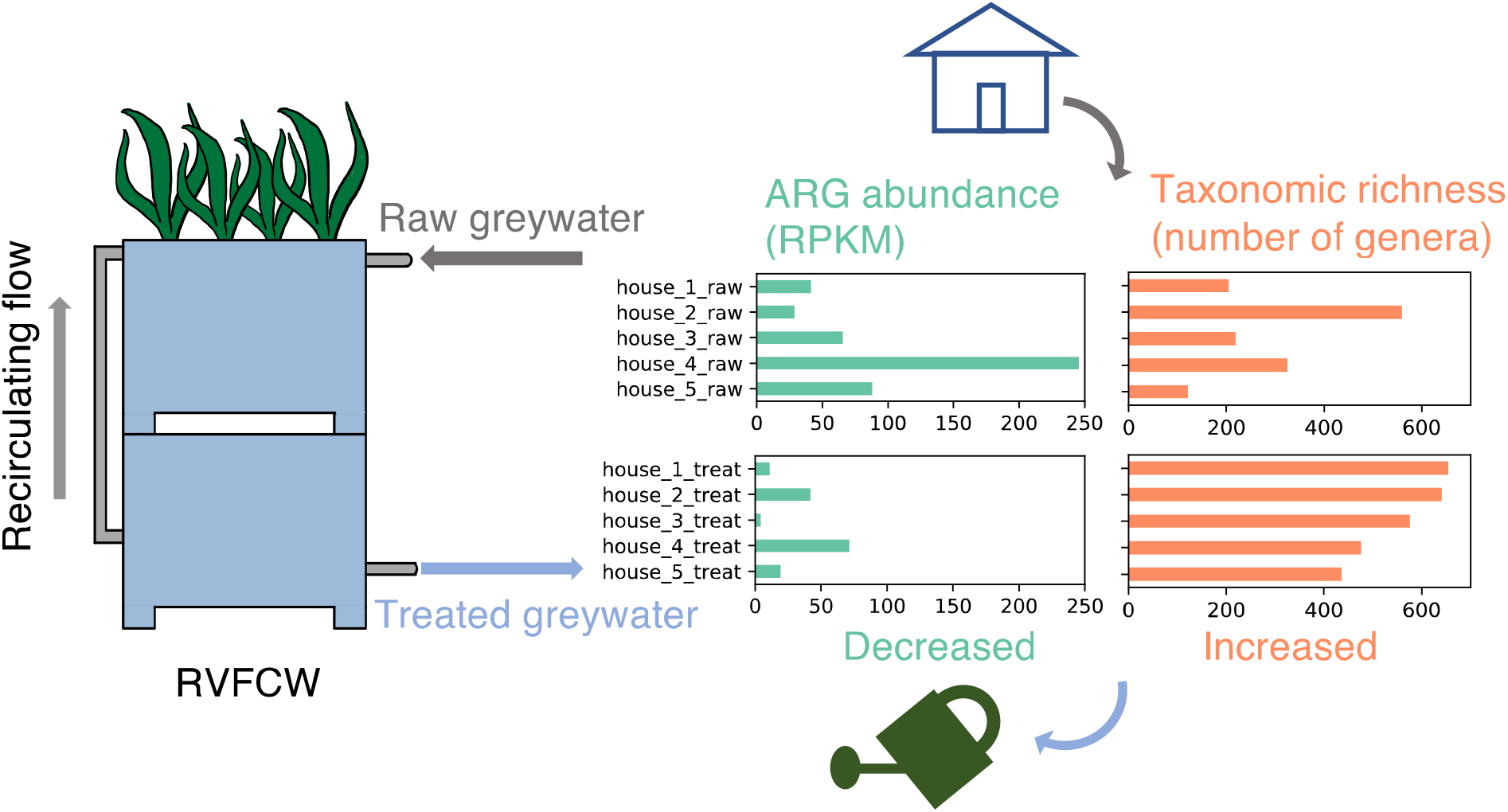

## Notes

### Competing Interest Statement

The authors have declared no competing interest.

