## Supporting Information for "Mitigation of Antimicrobial Resistance Genes in Greywater Treated at Household Level"

**Table S1.** Raw and decontaminated sequence numbers and average length for greywater samples.

| Sample | Raw sequence | | Decontaminated sequence | |
|---|---|---|---|---|
| | Total Sequence | Average Sequence Length | Total Sequence | Average Sequence Length |
| house_1_raw_R1 | 41542263 | 150 | 34364929 | 148.2 |
| house_1_raw_R2 | 41542263 | 150 | 34364929 | 147.6 |
| house_1_treat_R1 | 39393230 | 150 | 32050785 | 148.1 |
| house_1_treat_R2 | 39393230 | 150 | 32050785 | 147.3 |
| house_2_raw_R1 | 40411838 | 150 | 33641442 | 148.3 |
| house_2_raw_R2 | 40411838 | 150 | 33641442 | 147.4 |
| house_2_treat_R1 | 46908121 | 150 | 38434083 | 148.4 |
| house_2_treat_R2 | 46908121 | 150 | 38434083 | 147.4 |
| house_3_raw_R1 | 33239379 | 150 | 27736047 | 148.3 |
| house_3_raw_R2 | 33239379 | 150 | 27736047 | 147.6 |
| house_3_treat_R1 | 34582286 | 150 | 28408621 | 148.2 |
| house_3_treat_R2 | 34582286 | 150 | 28408621 | 147.5 |
| house_4_raw_R1 | 28053838 | 150 | 23416543 | 148.3 |
| house_4_raw_R2 | 28053838 | 150 | 23416543 | 147.7 |
| house_4_treat_R1 | 36739110 | 150 | 30347847 | 148.3 |
| house_4_treat_R2 | 36739110 | 150 | 30347847 | 147.4 |
| house_5_raw_R1 | 31486921 | 150 | 26152913 | 148.3 |
| house_5_raw_R2 | 31486921 | 150 | 26152913 | 147.7 |
| house_5_treat_R1 | 25580381 | 150 | 20646030 | 148.1 |
| house_5_treat_R2 | 25580381 | 150 | 20646030 | 147.4 |
| kit_control_1_R1 | 14890 | 150 | | |
| kit_control_1_R2 | 14890 | 150 | | |
| kit_control_2_R1 | 4704 | 150 | | |
| kit_control_2_R2 | 4704 | 150 | | |

**Table S2.** Metagenome assembly quality assessed by QUAST based on metaSPAdes output.

| | Number of contigs | | N50 | | Genome fraction (%) | |
|---|---|---|---|---|---|---|
| | Raw | Treated | Raw | Treated | Raw | Treated |
| House 1 | $5.97\times 10^4$ | $5.23\times 10^5$ | 4151 | 1410 | 23.9 | 1.1 |
| House 2 | $3.75\times 10^5$ | $4.86\times 10^5$ | 2124 | 1280 | 30.4 | 17.4 |
| House 3 | $8.24\times 10^4$ | $3.56\times 10^5$ | 2867 | 2006 | 28.7 | 10.7 |
| House 4 | $1.18\times 10^5$ | $2.64\times 10^5$ | 3185 | 1915 | 22.1 | 12.8 |
| House 5 | $4.57\times 10^4$ | $3.49\times 10^5$ | 5476 | 1065 | 28.6 | 12.5 |
| Average | $1.36\times 10^5$ | $3.96\times 10^5$ | 3561 | 1535 | 26.7 | 10.9 |
| p value[1] | 0.014* | | 0.043* | | 0.002* | |

[1] Two-tailed, paired two sample *t*-test.

* p value < 0.05.

**Table S3**. Relative abundance (%) of clinically important taxa among the top 50 species identified by MetaPhlAn 2.0.

| Clinical importance | Taxonomic classification | House 1 | | House 2 | | House 3 | | House 4 | | House 5 | |
|---|---|---|---|---|---|---|---|---|---|---|---|
| | | Raw | Treated | Raw | Treated | Raw | Treated | Raw | Treated | Raw | Treat |
| Certain species known human pathogen | *Pseudomonas* unclassified | 48.7 | 17.0 | 13.5 | 12.8 | 4.0 | 18.2 | 4.0 | 1.6 | 9.2 | 16.9 |
| | *Acinetobacter* unclassified | 0.005 | 5.0 | 5.9 | 67.5 | 3.8 | 41.6 | 0.05 | 2.6 | 9.1 | 7.1 |
| | *Arcobacter* unclassified | 48.2 | 0.02 | 5.4 | 1.8 | 0.01 | 2.0 | 0.2 | 5.4 | 4.7 | 0.2 |
| | *Aeromonas* unclassified | 0.02 | - | 2.3 | 0.4 | 8.8 | 4.9 | 0.2 | 0.05 | 0.9 | 1.0 |
| | *Comamonas* unclassified | 0.004 | 0.07 | 0.7 | 0.05 | 0.06 | 0.04 | 0.4 | - | 1.7 | 1.5 |
| | *Caulobacter* unclassified | 0.05 | 0.2 | 0.006 | 0.3 | 0.003 | 0.2 | 0.1 | 1.3 | - | 0.8 |
| | *Brevundimonas* unclassified | - | 0.2 | 0.2 | 0.06 | 0.07 | 0.9 | 0.02 | 0.2 | 0.02 | 0.4 |
| | *Sphingobium* unclassified | - | 0.1 | 0.1 | 0.2 | - | - | - | 0.7 | - | 0.7 |
| | *Paracoccus* unclassified | 0.01 | 0.3 | 0.5 | 0.04 | 0.03 | 0.2 | - | 0.1 | - | 0.4 |
| Common human pathogen | *Citrobacter freundii* | 0.4 | - | 1.3 | - | 0.006 | - | 0.07 | - | 0.07 | - |
| | *Klebsiella oxytoca* | - | - | - | - | 0.03 | - | 1.0 | - | 0.4 | 0.3 |
| | *Desulfovibrio desulfuricans* | 0.6 | - | 1.0 | - | 0.03 | - | 0.2 | - | - | - |
| Less common human pathogen, possible opportunistic infection in immunocompromised host | *Acinetobacter johnsonii* | - | 0.2 | 13.7 | 0.2 | 0.009 | 0.3 | 0 | 44.1 | - | - |
| | *Acinetobacter junii* | - | - | 6.2 | 2.4 | 0.9 | 0.2 | 0.2 | 25.3 | 3.7 | 0.6 |
| | *Arcobacter butzleri* | 0.09 | - | 0.2 | 0.02 | - | - | 0.002 | 0.08 | 14.1 | - |
| | *Laribacter hongkongensis* | 0.2 | - | 10.7 | - | - | - | - | - | - | - |
| | *Pseudomonas putida* | - | 0.2 | 0.04 | 0.08 | - | - | - | 0.01 | 9.4 | - |
| | *Azospira oryzae* | 0.02 | - | 1.6 | 0.004 | 0.0004 | - | 0.9 | 0.08 | 1.3 | 0.5 |
| | *Aeromonas caviae* | 0.006 | - | 0.5 | 0.1 | 1.9 | 0.7 | 0.02 | - | 0.8 | - |
| | *Pseudomonas alcaligenes* | - | - | - | - | 0.02 | - | - | - | 2.8 | - |
| | *Aeromonas hydrophila* | - | - | 1.4 | 0.6 | 0.001 | 0.2 | - | - | 0.1 | - |
| Non-pathogenic but potential carrier of ARG | *Parabacteroides* unclassified | 0.1 | - | 4.0 | - | - | - | 0.5 | - | 0.06 | - |

**Table S4.** Spearman's rank correlation between metaxa2 identified genus and RGI-bwt identified ARG alleles ($r \geq 0.6$, $p < 0.05$, excluding genus-ARG pair with only two sets of non-zero values among 10 greywater samples).

| Genus | ARG allele | Spearman's rank correlation (r) | p value |
|---|---|---|---|
| *Novispirillum* | *aac(3)-Ib*$^*$ | 1.000 | 0.002 |
| *Desulfobulbus* | *bla*$_{\text{AER-1}}$$^*$ | 1.000 | 0.007 |
| *Pelobacter* | | 1.000 | 0.017 |
| *Syntrophomonas* | | 1.000 | 0.000 |
| *Thiovirga* | | 1.000 | 0.047 |
| *Limnohabitans* | *adeF*$^*$ | 1.000 | 0.026 |
| *Bacteroides* | *mdtC* | 1.000 | 0.012 |
| *Novispirillum* | *sul2*$^*$ | 1.000 | 0.024 |
| *Macromonas* | *tetA*$^*$ | 1.000 | 0.046 |
| *Novispirillum* | | 1.000 | 0.024 |
| *Bacteroides* | *tetC* | 1.000 | 0.031 |
| *Citrobacter* | | 1.000 | 0.014 |
| *Laribacter* | | 1.000 | 0.003 |
| *Parabacteroides*$^*$ | | 1.000 | 0.012 |
| *Rhodocyclus* | | 1.000 | 0.010 |
| *Acidovorax*$^*$ | *adeF*$^*$ | 0.881 | 0.001 |
| *Brachymonas* | *bla*$_{\text{AER-1}}$$^*$ | 0.829 | 0.007 |
| *Desulforegula* | | 0.829 | 0.034 |
| *Propionivibrio* | | 0.829 | 0.002 |
| *Tolumonas*$^*$ | *sul2*$^*$ | 0.829 | 0.034 |
| | *tetA*$^*$ | 0.829 | 0.032 |
| | *aac(3)-Ib*$^*$ | 0.800 | 0.039 |
| *Dickeya* | *aph(6)-Id*$^*$ | 0.800 | 0.000 |
| *Thauera* | | 0.800 | 0.023 |
| *Tolumonas*$^*$ | | 0.800 | 0.012 |
| *Simplicispira* | *adeF*$^*$ | 0.800 | 0.027 |
| *Dickeya* | *tetA*$^*$ | 0.800 | 0.001 |
| *Cloacibacterium*$^*$ | *bla*$_{\text{CTX-M-126}}$$^*$ | 0.771 | 0.034 |
| *Azospira* | | 0.714 | 0.001 |
| *Methanosaeta* | *bla*$_{\text{AER-1}}$$^*$ | 0.700 | 0.036 |
| *Thauera* | *tetA*$^*$ | 0.679 | 0.005 |
| *Massilia* | *adeF*$^*$ | 0.667 | 0.028 |
| *Rhodocyclus* | *bla*$_{\text{AER-1}}$$^*$ | 0.600 | 0.006 |
| *Enhydrobacter* | *bla*$_{\text{CTX-M-126}}$$^*$ | 0.600 | 0.031 |

$^*$ Top 20 most abundant genera or top 10 most abundant ARG alleles.

**Table S5.** Top five ARG-associated species detected in greywater samples and ARGs found on the corresponding contigs.

| Top 5 ARG-associated species | | ARGs identified on contigs |
|---|---|---|
| **Raw greywater** | *P. aeruginosa* | *lcr-1, tetC, aph(3")-Ib, aph(6)-Id, bla*$_{OXA-10}$, *qnrVC1, aadA4, aadA7, arr-2, dfrA22, qacEdelta1* |
| | *E. coli* | *aph(3")-Ib, aph(6)-Id, aadA10, mef(B), qacEdelta1, sul1* |
| | *C. freundii* | *CpxA*, CRP, *Escherichia coli glpT* with mutation conferring resistance to fosfomycin, *Escherichia coli uhpT* with mutation conferring resistance to fosfomycin, H-NS, *emrB* |
| | *P. alcaligenes* | *rsmA* |
| | *K. pneumoniae* | *bla*$_{OXA-4}$, *cmlA4, mphE, msrE* |
| **Treated greywater** | *K. grimontii* | *bla*$_{OXA-10}$, *qacEdelta1, sul1, tetA* |
| | *A. baumannii* | *bla*$_{OXA-58}$, *mphE, msrE, tet(39)* |
| | *A. gyllenbergii* | *adeF* |
| | *E. coli* | *aph(3")-Ib, aph(6)-Id* |
| | *A. dispersus* | *ant(3")-IIc, adeF* |

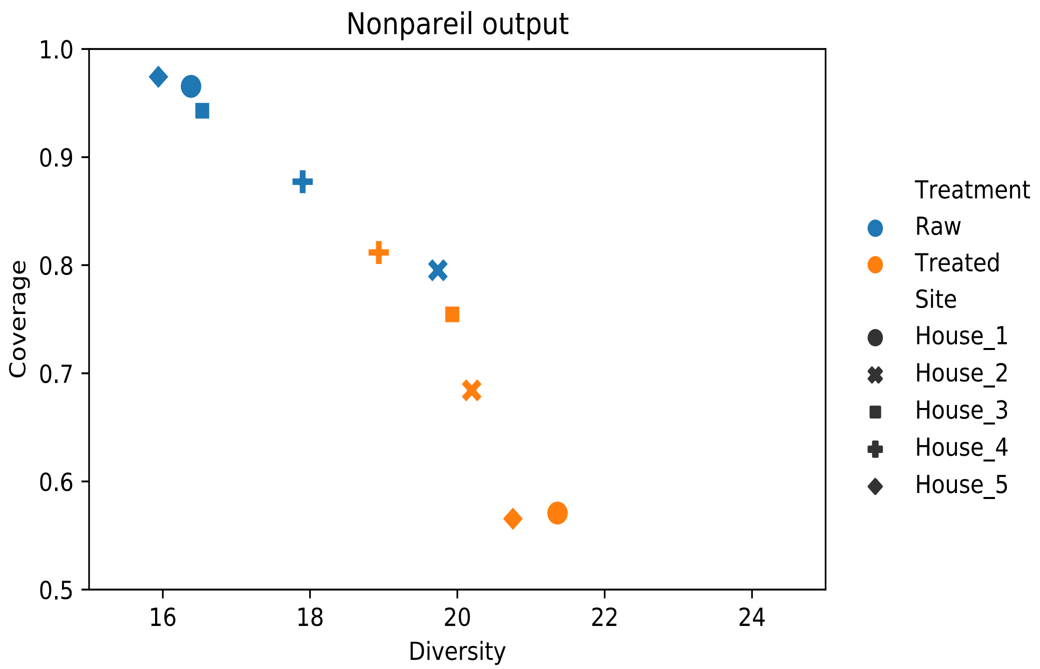


**Figure S1.** Nonpareil estimated metagenome coverage and diversity.

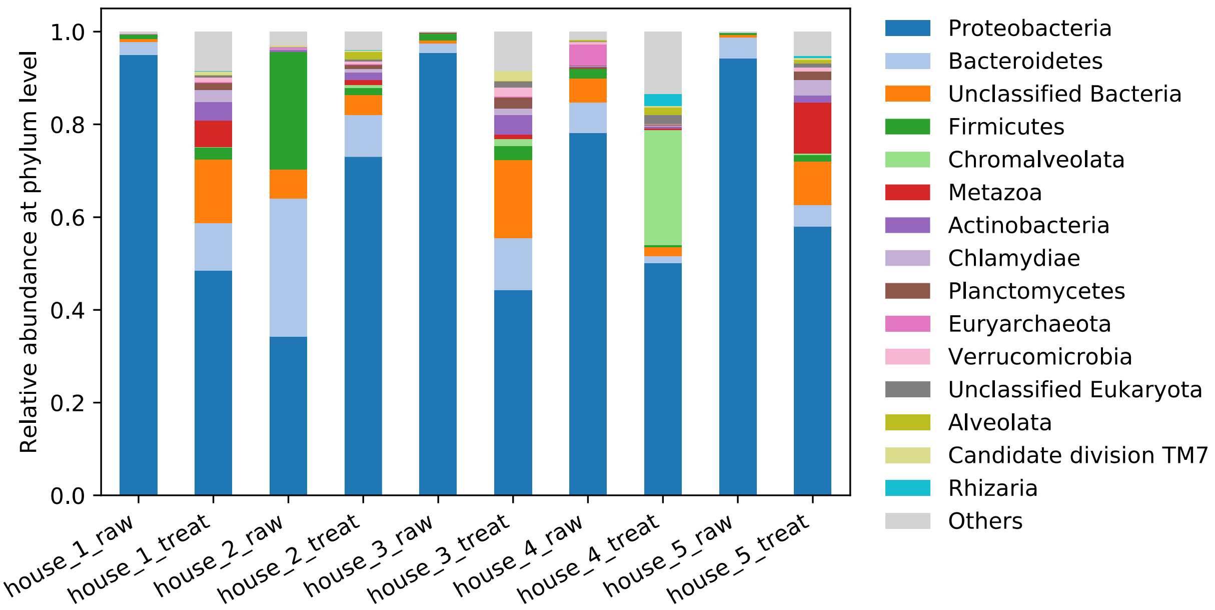


**Figure S2.** Community composition at phylum level identified by Metaxa2.

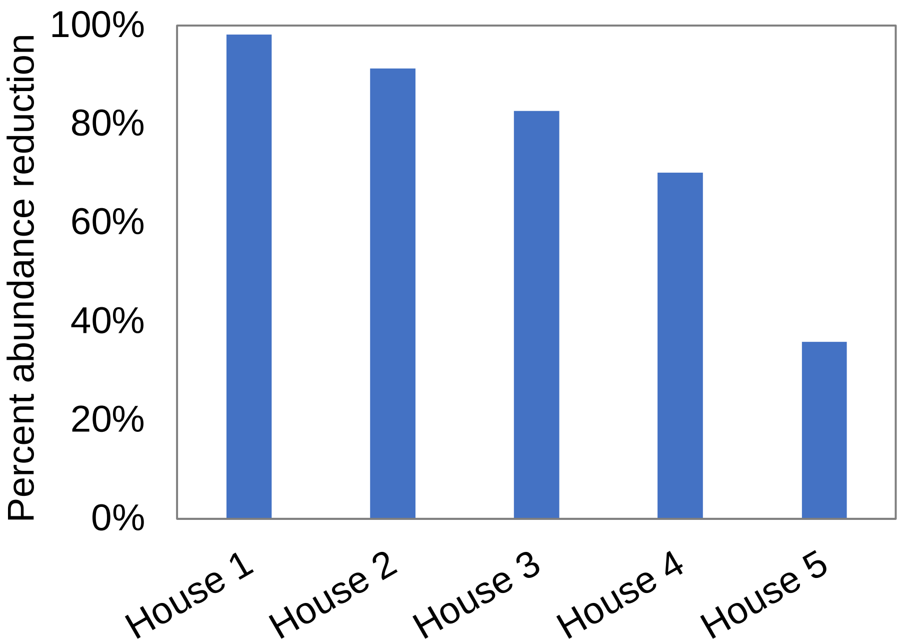


**Figure S3.** Percent abundance reduction for non-efflux resistance genes in the five households after treatment by the RVFCW facilities. Reduction was calculated from ARG relative abundance in RPKM.

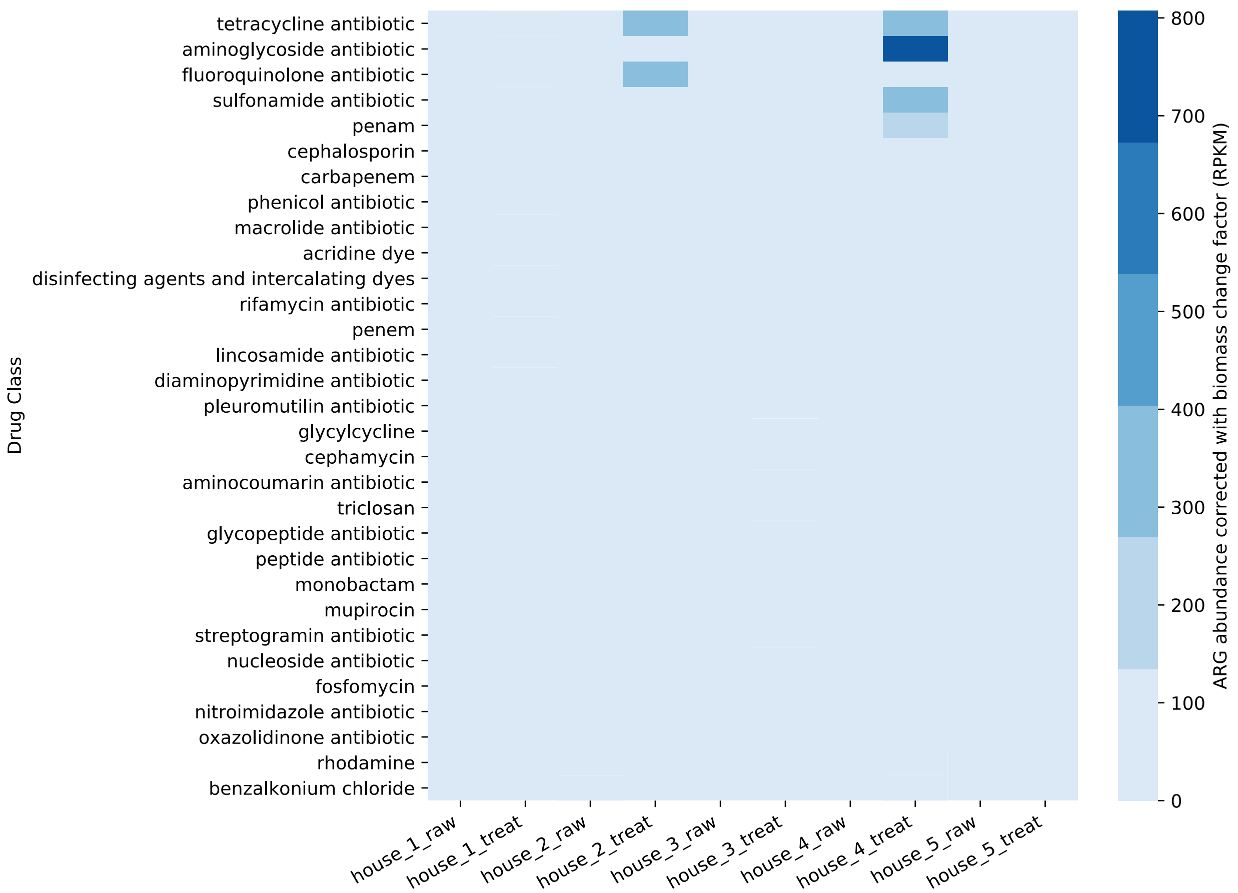


**Figure S4**. Abundance of antimicrobial resistance genes (RPKM) corrected with total biomass change factor after treatment by recirculating vertical flow constructed wetland reactors.

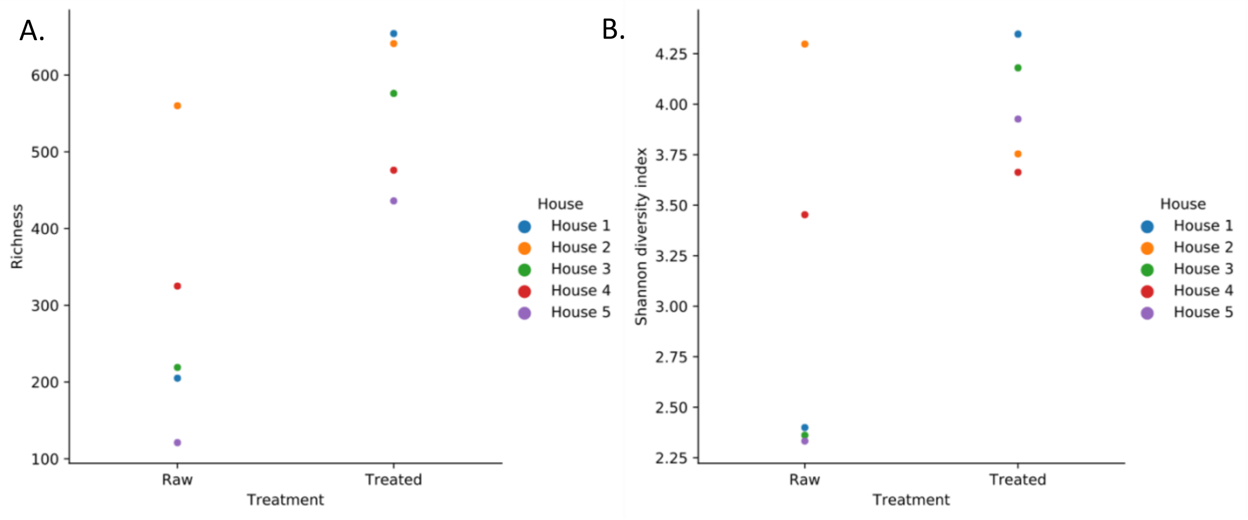


**Figure S5.** Taxonomic diversity expressed as genera richness (A) and Shannon diversity index (B) before and after treatment of greywater samples.

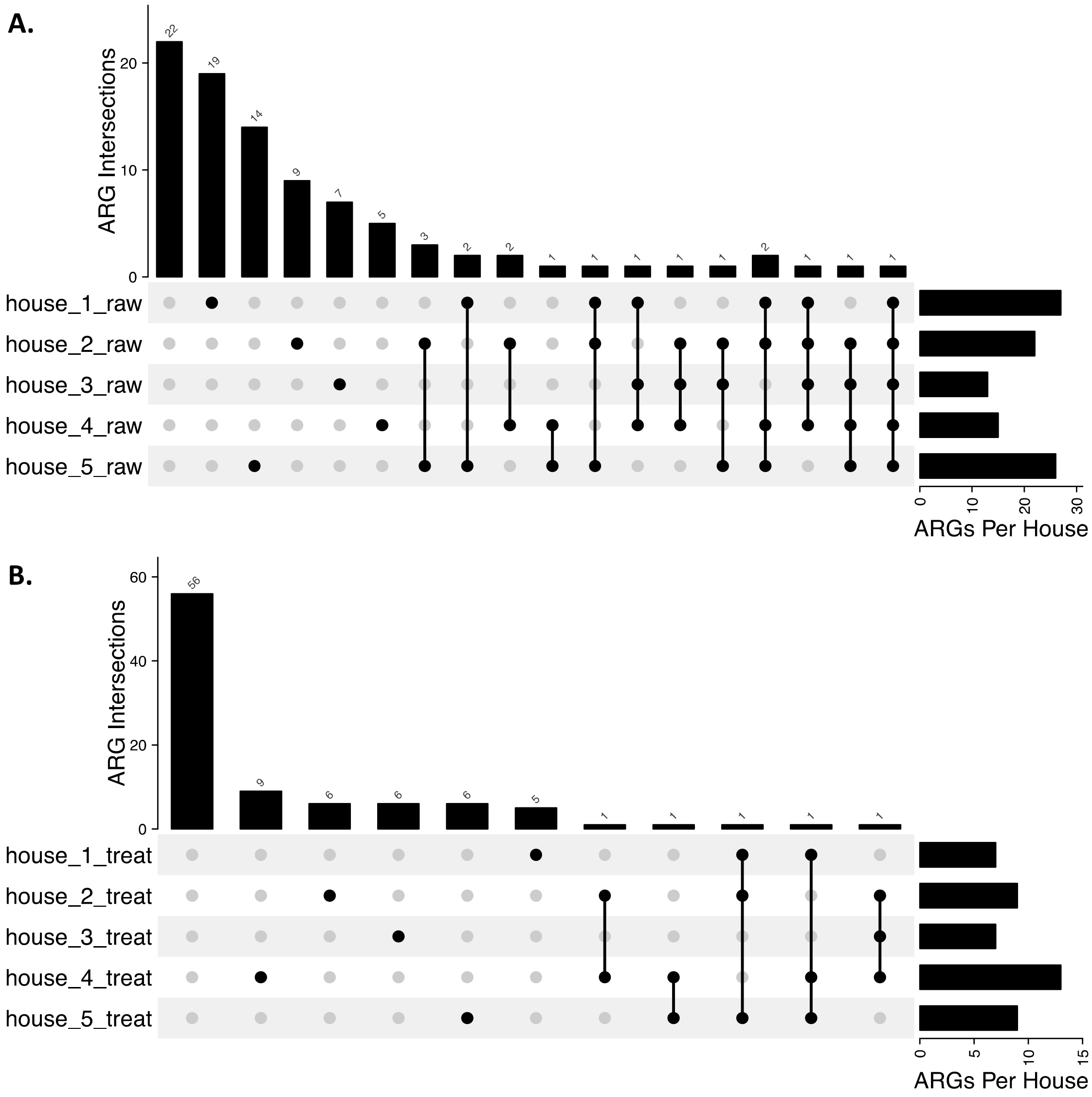


**Figure S6.** Number of shared and unique antimicrobial resistance genes (ARGs) identified from raw (A) and treated (B) greywater samples. First column in raw greywater plot represents ARG alleles appeared only in treated greywater samples, and *vice versa*. Results are based on short read-based analysis using RGI from CARD.

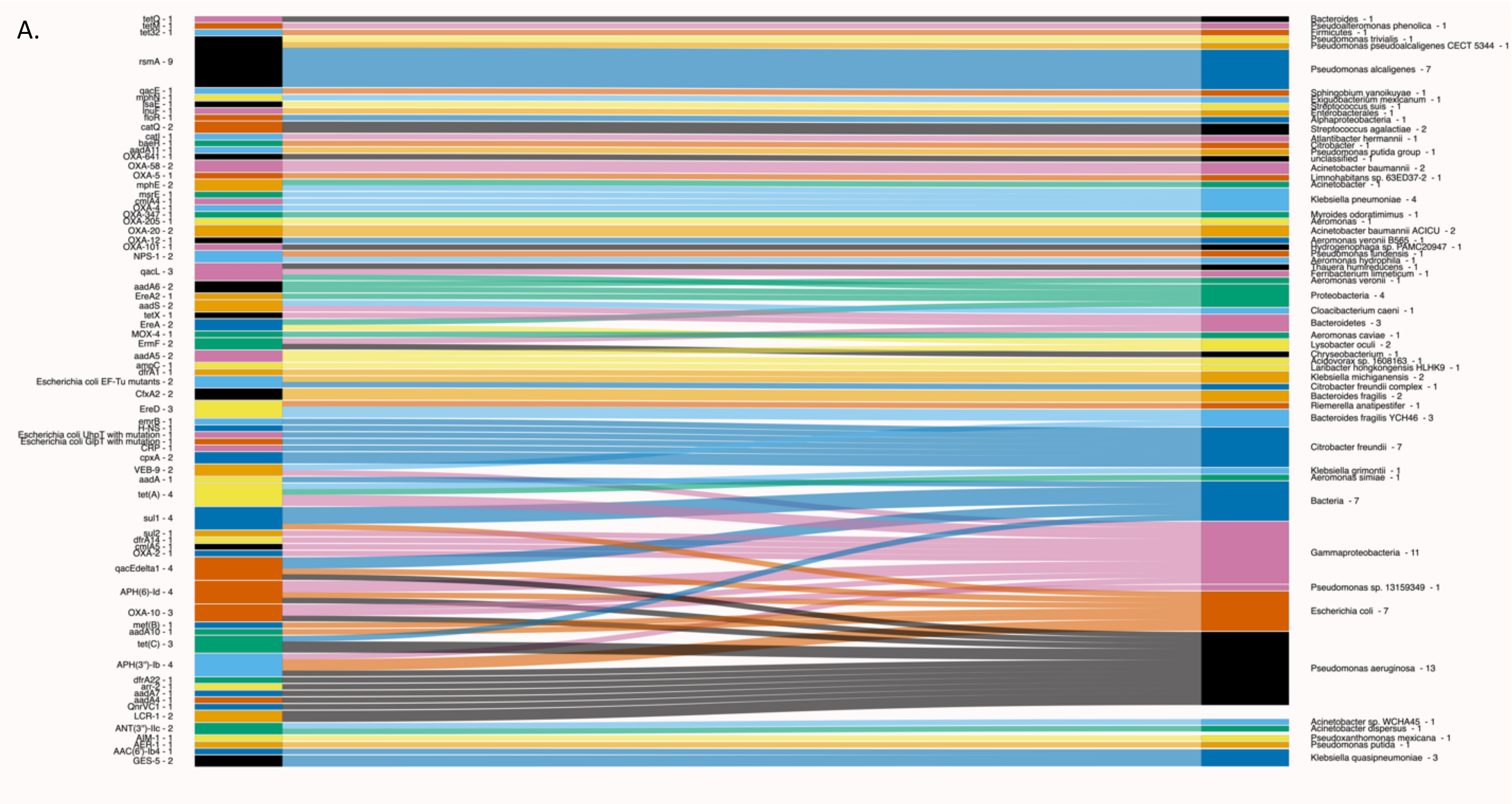
A.
tetQ - 1
tetM - 1
tet32 - 1
rsmA - 9
qacE - 1
mphN - 1
lsaE - 1
lnuF - 1
floR - 1
catQ - 2
catI - 1
baeR - 1
aadA11 - 1
OXA-641 - 1
OXA-58 - 2
OXA-5 - 1
mphE - 2
msrE - 1
cmlA4 - 1
OXA-4 - 1
OXA-347 - 1
OXA-205 - 1
OXA-20 - 2
OXA-12 - 1
OXA-101 - 1
NPS-1 - 2
qacL - 3
aadA6 - 2
EreA2 - 1
aadS - 2
tetX - 1
EreA - 2
MOX-4 - 1
ErmF - 2
aadA5 - 2
ampC - 1
dfrA1 - 1
Escherichia coli EF-Tu mutants - 2
CfxA2 - 2
EreD - 3
emrB - 1
H-NS - 1
Escherichia coli UhpT with mutation - 1
Escherichia coli GlpT with mutation - 1
CRP - 1
cpxA - 2
VEB-9 - 2
aadA - 1
tet(A) - 4
sul1 - 4
sul2 - 1
dfrA14 - 1
cmlA5 - 1
OXA-2 - 1
qacEdelta1 - 4
APH(6)-Id - 4
OXA-10 - 3
mef(B) - 1
aadA10 - 1
tet(C) - 3
APH(3'')-Ib - 4
dfrA22 - 1
arr-2 - 1
aadA7 - 1
aadA4 - 1
QnrVC1 - 1
LCR-1 - 2
ANT(3'')-IIc - 2
AIM-1 - 1
AER-1 - 1
AAC(6')-Ib4 - 1
GES-5 - 2
Bacteroides - 1
Pseudoalteromonas phenolica - 1
Firmicutes - 1
Pseudomonas trivialis - 1
Pseudomonas pseudoalcaligenes CECT 5344 - 1
Pseudomonas alcaligenes - 7
Sphingobium yanoikuyae - 1
Exiguobacterium mexicanum - 1
Streptococcus suis - 1
Enterobacterales - 1
Alphaproteobacteria - 1
Streptococcus agalactiae - 2
Atlantibacter hermannii - 1
Citrobacter - 1
Pseudomonas putida group - 1
unclassified - 1
Acinetobacter baumannii - 2
Limnohabitans sp. 63ED37-2 - 1
Acinetobacter - 1
Klebsiella pneumoniae - 4
Myroides odoratimimus - 1
Aeromonas - 1
Acinetobacter baumannii ACICU - 2
Aeromonas veronii B565 - 1
Hydrogenophaga sp. PAMC20947 - 1
Pseudomonas lundensis - 1
Aeromonas hydrophila - 1
Thauera humireducens - 1
Ferribacterium limneticum - 1
Aeromonas veronii - 1
Proteobacteria - 4
Cloacibacterium caeni - 1
Bacteroidetes - 3
Aeromonas caviae - 1
Lysobacter oculi - 2
Chryseobacterium - 1
Acidovorax sp. 1608163 - 1
Laribacter hongkongensis HLHK9 - 1
Klebsiella michiganensis - 2
Citrobacter freundii complex - 1
Bacteroides fragilis - 2
Riemerella anatipestifer - 1
Bacteroides fragilis YCH46 - 3
Citrobacter freundii - 7
Klebsiella grimontii - 1
Aeromonas simiae - 1
Bacteria - 7
Gammaproteobacteria - 11
Pseudomonas sp. 13159349 - 1
Escherichia coli - 7
Pseudomonas aeruginosa - 13
Acinetobacter sp. WCHA45 - 1
Acinetobacter dispersus - 1
Pseudoxanthomonas mexicana - 1
Pseudomonas putida - 1
Klebsiella quasipneumoniae - 3

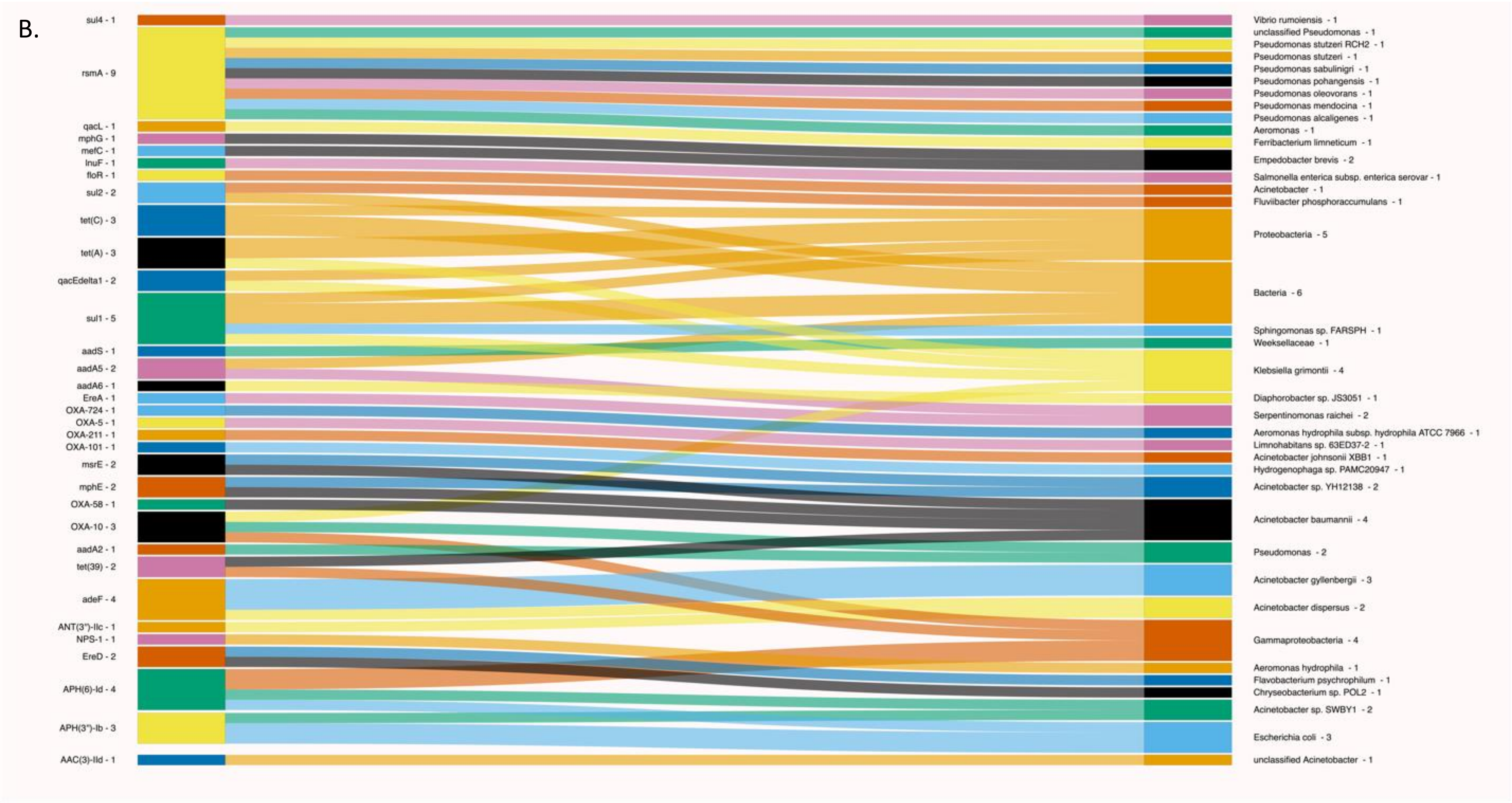


**Figure S7.** Association of ARGs and taxa in raw (A) and treated (B) greywater samples based on contig-level analysis. Occurrence (count) is listed after each ARG or taxa name.

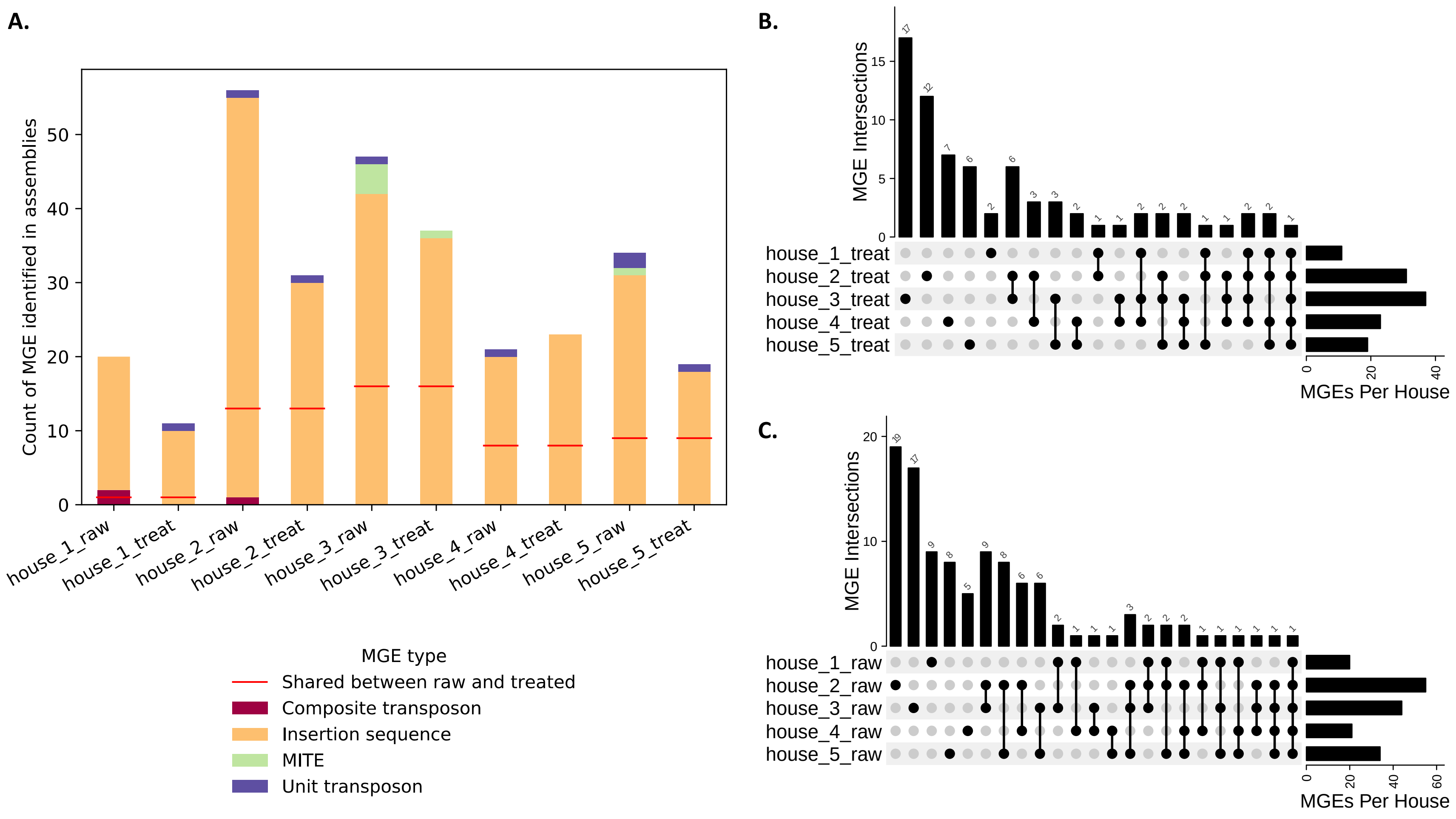


**Figure S8.** Occurrence (count) of mobile genetic elements (MGE) in greywater sample assemblies (A). Number of shared and unique MGEs identified on assemblies in raw (B) and treated (C) greywater samples.

### Quantitative Polymerase Chain Reaction Quality Control Information

Samples were transported to the laboratory within 4 hours of sampling and were filtered immediately. The filters were frozen at -20°C till DNA extraction, which happened within 4 working days. Extracted DNA were stored at -20°C until used for qPCR analysis. qPCR was performed with 20 µL reaction volumes including 1µL sample DNA. Primer set 341F/518R (5'-CCTACGGGAGGCAGCAG-3'/5'- ATTACCGCGGCTGCTGG-3') targeting the 16S rRNA gene was used for total bacteria quantification. Technical replicates (n=3) were measured for all samples. CFX96 Touch Real-Time PCR Detection is the manufacturer of the qPCR machine. The standard curve is displayed in Figure S9, the efficiency is 108.3%. The complete qPCR protocol is displayed in Figure S10.

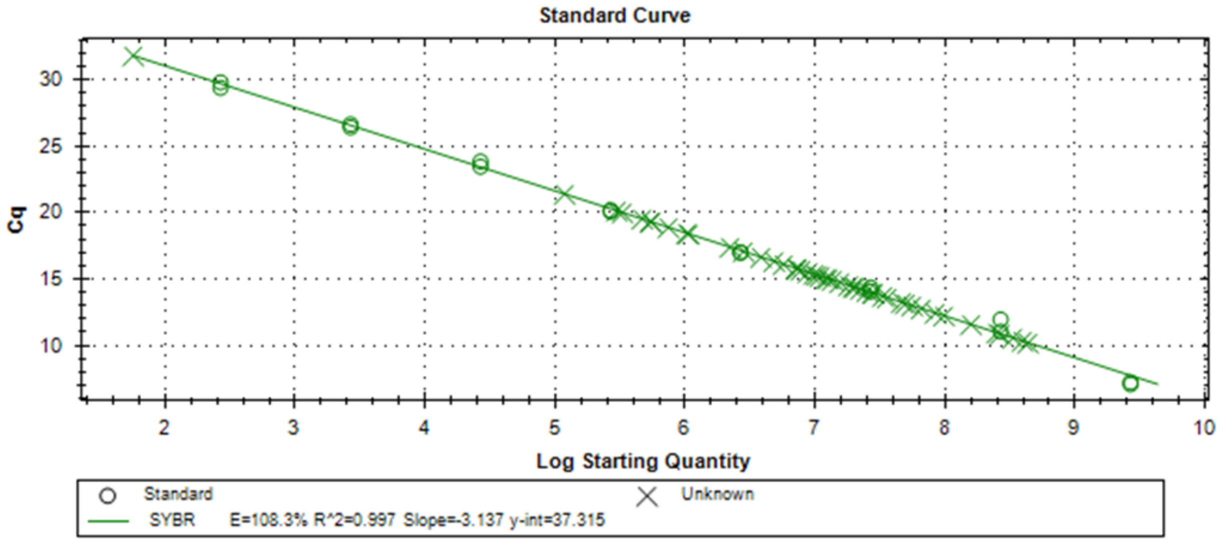


**Figure S9.** Standard curve of qPCR primer pair 341F/518R. Circular points are the standards and cross points are the samples.

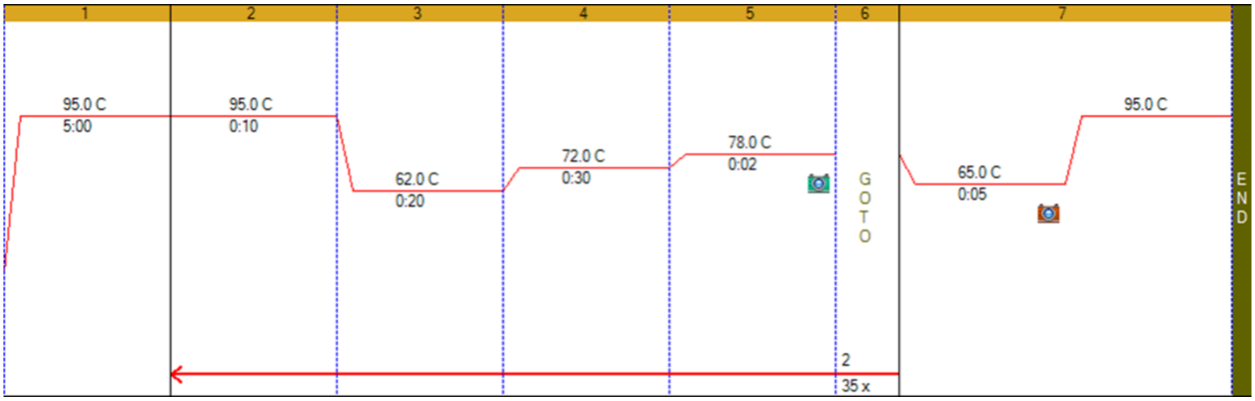


**Figure 10.** Protocol for qPCR reaction using primer pair 341F/518R.
